# S-Adenosyl-D-methionine as a Non-Physiological Substrate for a Wide Range of SAM-Dependent Enzymes

**DOI:** 10.64898/2026.04.20.719600

**Authors:** Philipp Germer, Lukas Gericke, Lars-Hendrik Koeppl, Ziruo Zou, Emely Jockmann, Marco Kuge, Katrin Zoller, Hannah Herrmann, Ronja Fuderer, Michael K. F. Mohr, Agnes Bartels, Göksu Oral, Peer Lukat, Gunhild Layer, Michael Müller, Wulf Blankenfeldt, Lena Barra, Jennifer N. Andexer

## Abstract

The ability of SAM-dependent enzymes to accept *S*-adenosyl-D-methionine [D-SAM, (*S*_S_,*R*_Cα_)-SAM] instead of the native cofactor *S*-adenosyl-L-methionine [L-SAM, (*S*_S_,*S*_Cα_)-SAM] remains largely unexplored. Challenging the stereochemical preference of SAM-dependent enzymes, we investigated the ability of different enzyme classes to accept D-SAM. Contrary to common assumptions, the tested *N*- and *O*-methyl transferases (MTs), as well as one of the examined *C*-MTs accepted D-SAM. Docking studies suggest that acceptance of D-SAM by *C*-MTs may be influenced by the angle between the transferable methyl group of SAM and the nucleophilic carbon of the substrate, along with enzyme and substrate flexibility. In addition to conventional MTs, the radical SAM glutamine *C*-MT QCMT showed low but detectable methylation activity with D-SAM. Furthermore, the azetidine-2-carboxylic acid synthase AzeJ not only uses D-SAM but also incorporates the stereocentre of D-methionine into the cyclic amino acid product. The pyridoxal 5′-phosphate (PLP)-dependent enzyme 1-aminocyclopropyl-1-carboxylic acid synthase (ACCS) also showed detectable turnover with D-SAM. These findings broaden the understanding of enzyme stereoselectivity, provide an overview of D-SAM-utilising enzymes, and identify first enzyme systems that may serve as starting points for engineering efforts aimed at shifting cofactor preference towards D-SAM.

## Introduction

*S*-Adenosyl-L-methionine [L-SAM, (*S*_S_,*S*_Cα_)-SAM] is an ubiquitous enzyme cofactor and a key player in one-carbon transfer,^[1]^ playing a central role in biomolecular methylation and epigenetic regulation.^[2]^ Beyond these functions, SAM participates in a broad array of biochemical transformations that proceed through distinct chemical mechanisms.^[3]^ Conventional SAM-dependent methyltransferases (MTs) typically catalyse S_N_2-type methyl transfer reactions at the sulfonium centre of SAM. Other SAM-dependent transformations include aminopropylation during polyamine biosynthesis, halogenation reactions catalysed by SAM-dependent halogenases, and radical-mediated reactions catalysed by the radical SAM enzyme superfamily.^[4]^ Members of this superfamily use a [4Fe-4S]-cluster to produce a 5′-deoxyadenosyl radical from SAM to initiate radical chemistry, enabling a wide variety of synthetically challenging transformations.^[5]^ Notably, some of the radical SAM enzymes, such as the cobalamin-dependent MT TsrM, perform methylation through alternative non-radical mechanisms.^[6]^ In addition, the cofactor is involved in many other reactions. Examples include the production of ethylene via pyridoxal 5′-phosphate (PLP)-dependent conversion of SAM to 1-aminocyclopropyl-1-carboxylic acid (ACC), catalysed by ACC synthase (ACCS) This reaction proceeds via formation of an external aldimine between the α-amino group of SAM and the PLP cofactor. Another example of a SAM-dependent enzyme catalysing a non-methyl transfer reaction is the azetidine-2-carboxylic acid (AZC) synthase AzeJ, which is related to the class I MTs on a sequence basis. In both cases, the Met-derived side chain of SAM undergoes cyclisation, yielding ACC in the ACCS-catalysed reaction and AZC in the AzeJ-catalysed reaction.^[4,7]^

The chemical structure of SAM was first elucidated by Cantoni in 1952.^[8]^ The cofactor contains an adenosyl unit connected to a Met moiety *via* a sulfonium. The resulting molecule contains six stereocentres: four in the ribose moiety of the adenosyl unit, one at the sulfonium centre, and one at the α-carbon of the Met-derived side chain. Enzymatically, SAM is synthesised from L-methionine (L-Met) and adenosine-5′-triphosphate (ATP) by methionine adenosyltransferases (MATs, Figure 1), with retention of the Cα-configuration, generating L-SAM.^1^ Interestingly, some MATs have been reported to accept D-methionine (D-Met) as a substrate, yielding D-SAM, the Cα-epimer of L-SAM, a property already described in the 1970s for the MAT from *Candida utilis* (syn. *Cyberlindnera jadinii*).^[9,10]^

**Figure 1.**
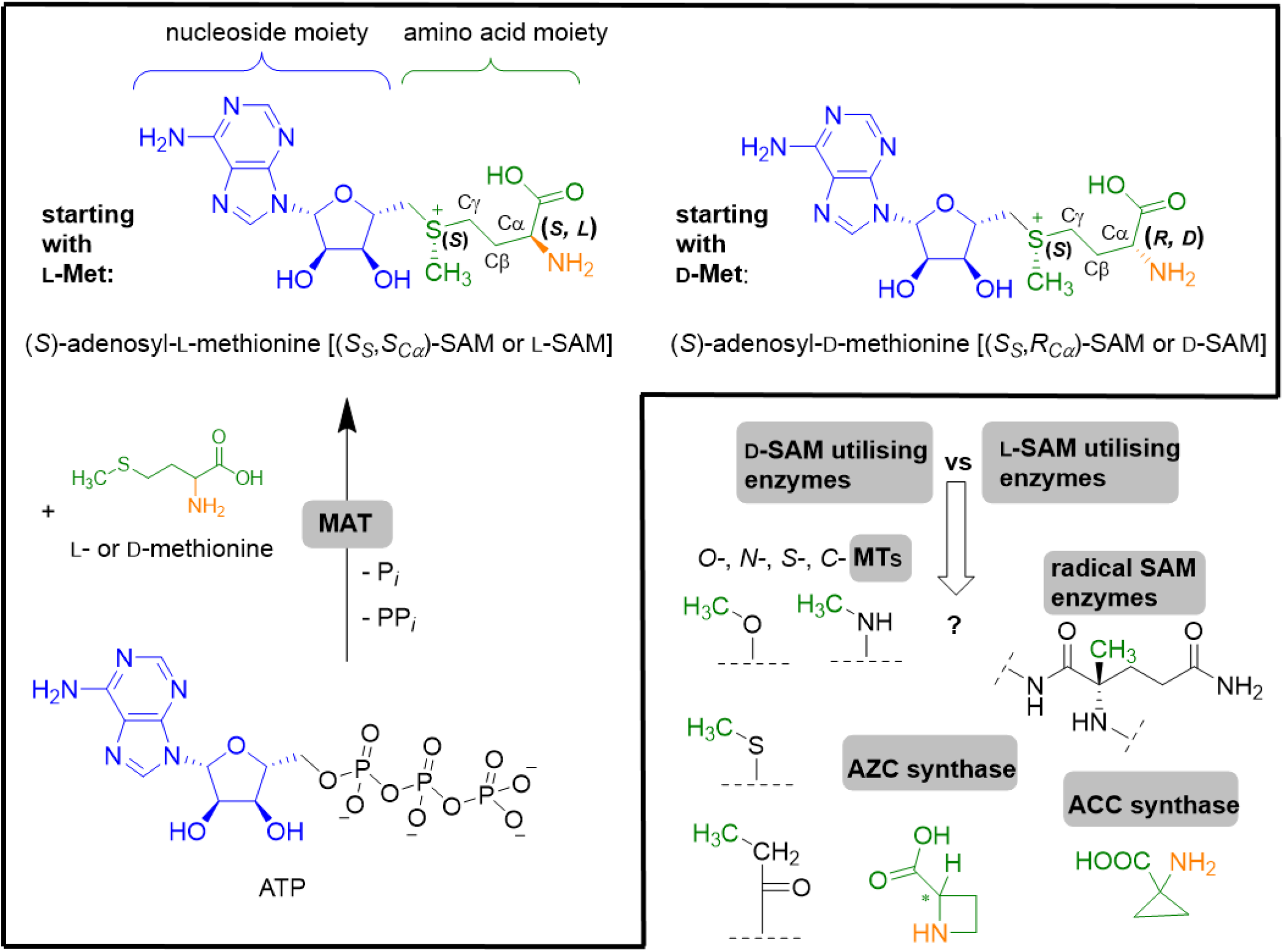
Methionine adenosyltransferase (MAT)-catalysed formation of the two diastereomers L-SAM and D-SAM starting from adenosine-5′-triphosphate (ATP) and L- or D-Met. In this work, we investigate the acceptance of D-SAM by diverse SAM-utilising enzymes including classical methyltransferases (MTs), radical SAM enzymes, 1-aminocyclopropyl-1-carboxylic acid synthase (ACCS) and azetidine-2-carboxylic acid (AZC) synthase.

Although the biological importance of L-SAM is well established, the potential for stereochemical variation at the α-amino acid position remains underexplored. Reports are limited to a few small molecule MTs, including *O*-, *N*-, and *S*-selective enzymes, which were either inhibited by D-SAM or showed decreased activity with D-SAM in comparison to L-SAM.^[10,11]^ The consequences of stereochemical inversion can be significant, particularly regarding cofactor recognition, catalytic performance and regulation. Such altered recognition may be desirable for biocatalytic applications, for example if an enzyme tolerates or selectively utilises D-SAM or related analogues without inhibition. This concept opens the possibility of creating orthogonal enzymatic systems, in which the use of D-Met or racemic Met can lead to the formation of D-SAM. Depending on whether the methyl group or the Met-derived side chain is transferred, D-SAM could then give rise to the same or different products as L-SAM, with the outcome further influenced by altered cofactor binding and reactivity. Recent work on SAM-dependent C-MTs has further shown that chemoselectivity can depend strongly on substrate binding geometry and precise positioning relative to SAM.^[12]^ Moreover, enzymes using L-SAM are highly diverse, reflecting the cofactor’s multifunctional nature, but also highlighting nature’s ability to recognise the cofactor through different structural binding modes. This raises the expectation that different SAM-dependent enzymes might be able to accept and use D-SAM in individual manners. Elucidation of the underlying mechanisms could give new insights into the control of stereoselectivity in SAM-dependent enzymes.

With this work, we set out to investigate the use of D-SAM by a panel of SAM-dependent enzymes (Figure 1). Firstly, we focused on SAM-dependent MTs with different substrates and chemoselectivities (*N*-, *O*-, and *C*-MTs) and then extended our screening towards radical SAM enzymes and Met side chain modifying enzymes, specifically AZC synthase and ACC synthase.^[13,14,15,16]^

## Results and Discussion

To investigate utilisation of D-SAM by SAM-dependent enzymes, a procedure had to be established to generate the desired SAM diastereomer. For this purpose, the MATs from *Thermococcus kodakarensis* (*Tk*MAT) and from *Ureaplasma urealyticum* (*Uu*MAT), which can generate D-SAM from D-Met and ATP, were used in an *in situ* cofactor supply cascade (see Figure 1, manuscript in preparation). Depending on the SAM-dependent enzyme used, turnover also generates characteristic byproducts: *S*-adenosylhomocysteine (SAH) for MTs, 5′-methylthioadenosine (MTA) for enzymes using the Met side chain, or 5′-deoxyadenosine (DOA) for radical SAM enzymes. In reactions producing SAH or MTA, MTA/SAH nucleosidase from *Escherichia coli* (*Ec*MTAN) was added to prevent byproduct accumulation.

This setup resulted in a three-enzyme cascade (MAT, SAM-dependent enzyme, MTAN), which was performed starting with D- and L-Met, forming the respective SAM diastereomer *in situ*. For radical SAM enzymes, *Ec*MTAN was omitted to allow analysis of the characteristic byproducts. In selected experiments, purified D-SAM and L-SAM were directly added to the enzyme of interest.

### Conventional *N*-, *O*-, and *C*-Methyltransferases

To assess D-SAM compatibility across different methylation chemistries, we selected representative SAM-dependent MTs covering *N*-, *O*-, and *C*-methylation. For *N*- and *O*-methylation, we chose the plant enzymes anthranilate *N*-MT from *Ruta graveolens* (*Rg*ANMT) and caffeate *O*-MT from *Prunus persica* (*Pp*CaOMT), two related small-molecule MTs. Both enzymes were previously shown to accept the non-natural substrate 2-amino-4-nitrophenol (2,4-ANP) when L-SAM is supplied.^[15]^ While *Rg*ANMT catalyses *N*-methylation resulting in the production of *N*-methyl-2,4-ANP, *Pp*CaOMT catalyses *O*-methylation to yield 2-methoxy-5-nitroanilline (2,5-MNA). For both MTs tested in the three-enzyme cascade, a product peak corresponding to the respective methylated product was observed when starting from D-Met. Comparison with assays starting from L-Met showed that product formation was generally slower when starting from D-Met. For *Pp*CaOMT, this was also reflected in a lower conversion under the tested conditions. For *Rg*ANMT, more than 99% of 2,4-ANP was converted to *N*-methyl-2,4-ANP after 24 hours of incubation with D-Met. *Pp*CaOMT reached about 92% *O*-methylation after 30 hours, despite the extended incubation time. In both cases, incubation with L-Met and thus L-SAM resulted in full conversion (over 99%). Due to the *in situ* production of SAM, it remains unclear whether these differences originate from slower D-SAM formation by MAT, from reduced catalytic efficiency of the MTs utilising D-SAM, or altered byproduct effects within the cascade, such as differences in SAH-mediated inhibition or turnover by MTAN.

The observed tolerance of the diastereomeric cofactor may reflect a less restrictive cofactor-binding site in *Rg*ANMT and *Pp*CaOMT, which appears to provide sufficient space and interactions to accommodate both SAM diastereomers.^[16]^ Despite slower product formation, both enzymes achieved high conversions with D-SAM, indicating that the stereochemical inversion at the α-carbon of SAM is tolerated by these enzymes. Since D-SAM has rarely been described as an active cofactor in enzymatic methylation reactions, our findings indicate that the stereochemical requirements of SAM-dependent MTs may be more flexible than previously anticipated. The high conversions suggest that the MAT-catalysed formation of D-SAM from D-Met is feasible and not a major limitation under the tested conditions. However, it remains possible that reduced MAT activity contributes to the slower overall methylation observed with D-Met as starting material.

Encouraged by these results, we investigated the compatibility of D-SAM with *C*-MTs. To cover two mechanistically distinct types of *C*-methylation, we selected CouO from *Streptomyces rishiriensis*, which acts on aromatic substrates such as 4,5,7-trihydroxy-3-phenylcoumarin, and SgvM from *Streptomyces griseoviridis*, a Zn^2+^-dependent enzyme modifying α-keto acids.^[17]^ Here, the nucleophile, a C-atom, requires more activation than N- and O-nucleophiles in *Pp*CaOMT’s and *Rg*ANMT’s reactions. Hence, correct substrate binding is more important.^[18,19]^ While CouO requires a near-perpendicular alignment between the aromatic substrate and the methyl group to enable electrophilic aromatic substitution, SgvM catalyses a *C*-methylation reaction on α-keto acids, which does not involve an aromatic π-system. Instead, efficient catalysis primarily depends on a favourable S_N_2-like geometry for nucleophilic substitution at the methyl group. Both enzymes were evaluated in the three-enzyme cascade as described before. When assayed with 4-methyl-2-oxovalerate, SgvM accepted both L- and D-SAM, although with lower product formation starting from D-Met, under the tested cascade conditions: Over 98% conversion was observed starting with L-Met after 33 hours, whereas around 29% product was observed starting from D-Met after the same incubation time (Figure S8). In contrast, CouO efficiently methylated 4,5,7-trihydroxy-3-phenylcoumarin in the presence of L-SAM (80% conversion after 24 hours) but showed no activity with D-SAM (Figure S6). These results suggest that D-SAM compatibility is not universal among *C*-MTs and may depend on specific structural and mechanistic features (Figure 2).

**Figure 2.**
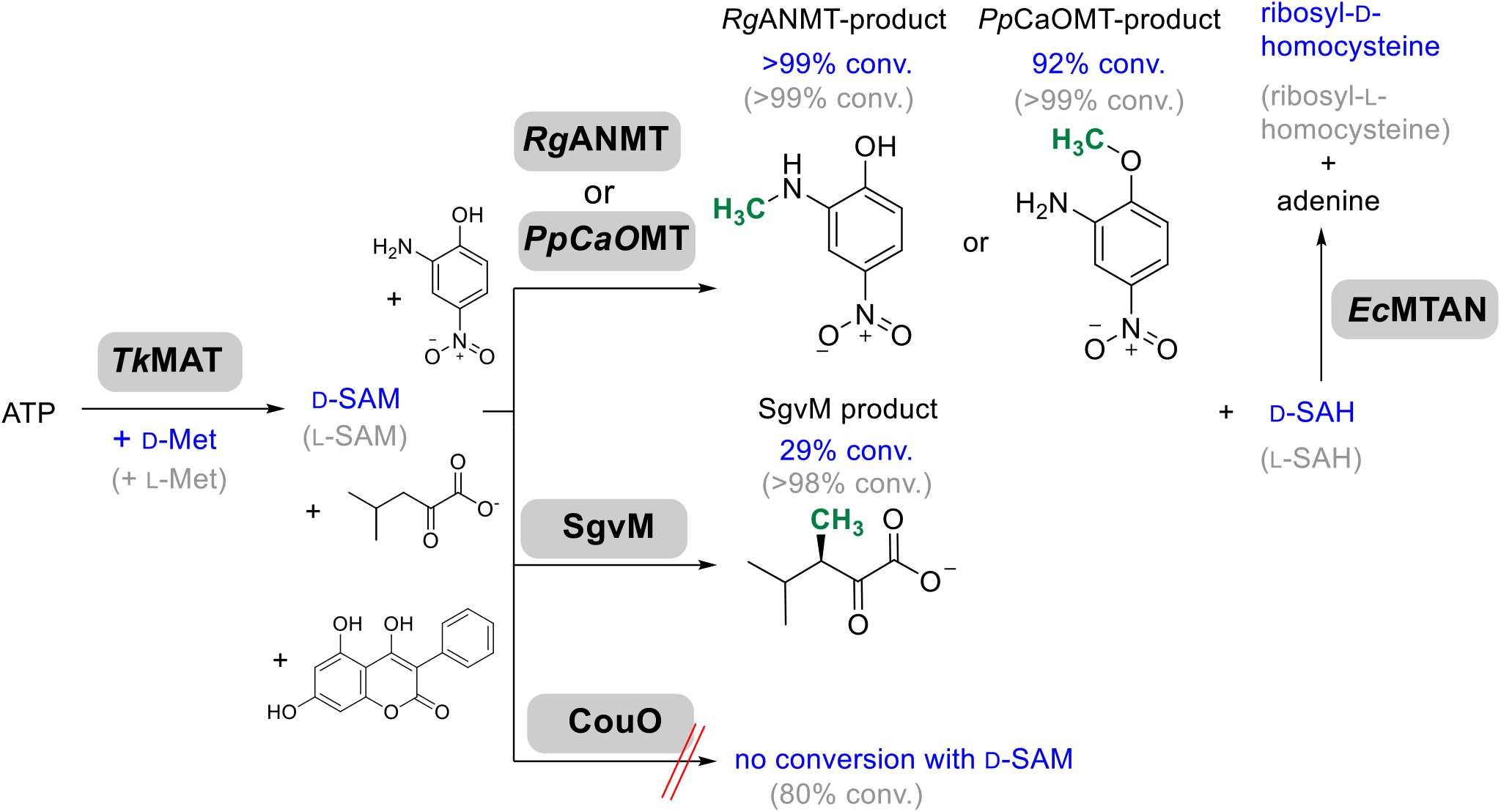
Three-enzyme cascade starting from ATP and D-Met to synthesise D-SAM with *Tk*MAT, followed by one of the MTs for methylation of its substrate and the generation of SAH, which is then degraded by *Ec*MTAN. *Rg*ANMT was used for *N*-methylation of 2-amino-4-nitrophenol (2,4-ANP), and *Pp*CaOMT was used for *O*-methylation of 2,4-ANP. The *C*-MT SgvM was used for methylation of 4-methyl-2-oxovalerate or 2-oxovalerate and *C*-MT CouO was tested with 4,5,7-trihydroxy-3-phenylcoumarin. Conversions with D-SAM are shown in blue, while conversions with L-SAM are shown in grey. Transferred methyl groups are highlighted green in the methylated products.

For CouO, docking studies using the experimentally tested substrate 4,5,7-trihydroxy-3-phenylcoumarin, D-SAM, and L-SAM were conducted based on the available crystal structure in complex with SAH (PDB: 5M58).^[20]^ This suggests that D-SAM adopts a binding orientation comparable to L-SAM. While the adenosyl moieties are largely superimposable, the amino acid moiety of D-SAM exhibits a slightly altered orientation. This results in a deviation of the transferable methyl group relative to the nucleophilic carbon of the aromatic substrate. The angle between the aromatic plane of the substrate and the electrophilic methyl group of L-SAM is approximately 93.1° (Figure S18 C), closely matching the near-perpendicular orientation expected to facilitate methyl transfer. In comparison, this angle increases to 105.4° with D-SAM (Figure S18 D), deviating from the optimal arrangement. Additionally, the angle formed between the nucleophilic carbon of the substrate, the methyl group, and the sulfonium sulfur is estimated to be 162.0° for L-SAM (Figure S18 A), approaching the near-linear geometry favourable for S_N_2-like methyl transfer, but decreases to 132.3° for D-SAM (Figure S18 B). In the D-SAM model, the methyl group is no longer positioned favourably relative to the nucleophilic carbon of the substrate. The substrate itself is stabilised by interactions with the side chains of R116, H117, H120, and R121 within the active site of CouO. Given the rigid, polyaromatic structure of 4,5,7-trihydroxy-3-phenylcoumarin, together with the amino acid interactions that anchor and orient it on both sides, the substrate-cofactor pair likely lacks the spatial freedom needed to compensate for the misalignment relative to the transferable methyl group of D-SAM. In addition, the guanidinium side chain of R116 forms a stabilising interaction with the carboxylate of L-SAM (Figure 3A). In the D-SAM complex (Figure 3B), the side chain of R116 may restrict the necessary positioning of the cofactor by repulsion of the positively charged amino group. Instead, the carboxyl group of D-SAM is reoriented and seems to be stabilised by Y25. Consequently, the orbital overlap required for electrophilic aromatic substitution is likely less favourable, which may prevent catalysis. Hence, for CouO, only the biogenic L-SAM supports efficient methyl transfer, whereas potential binding of D-SAM likely results in a catalytically incompetent enzyme-cofactor complex.

**Figure 3.**
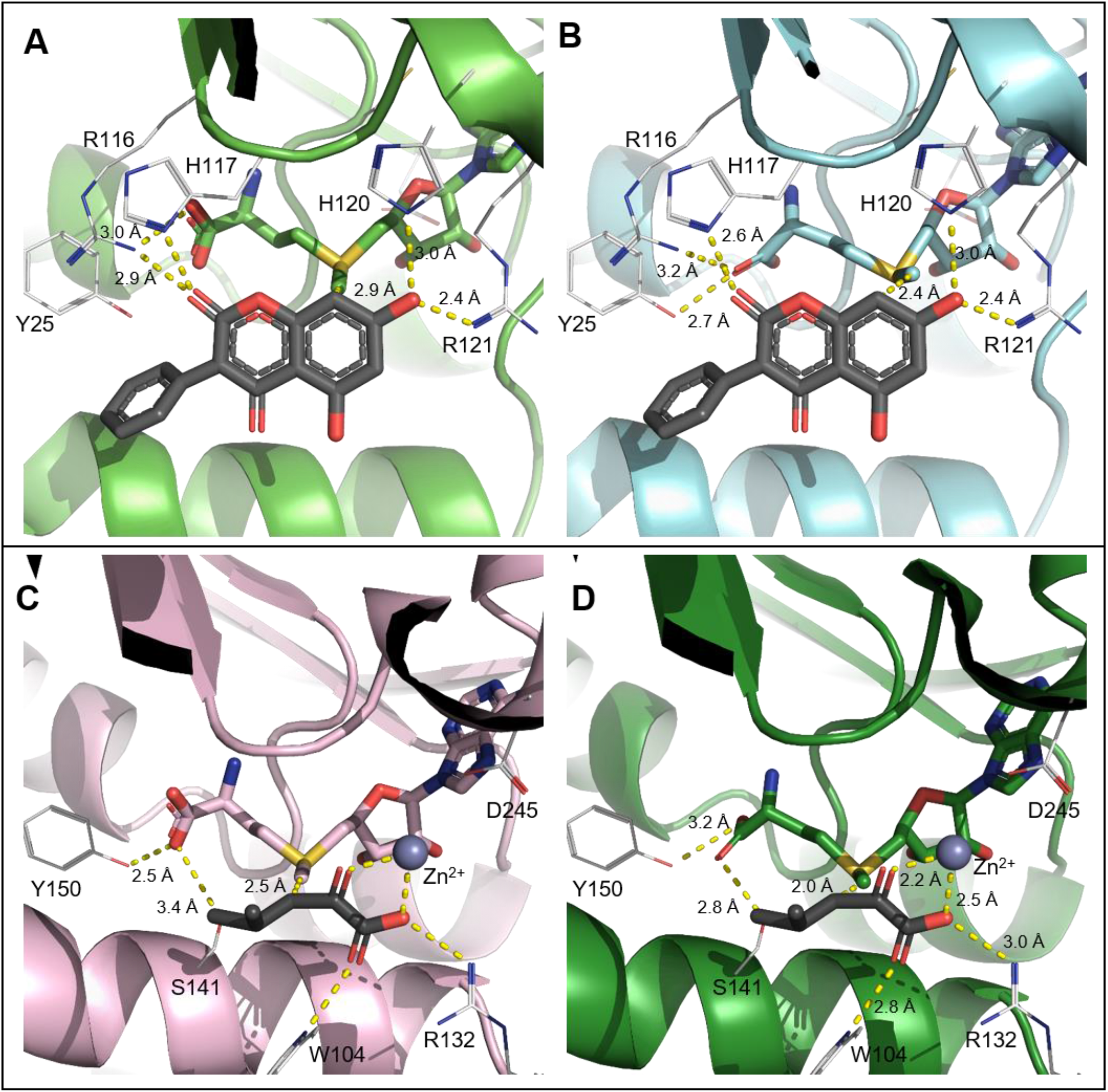
Docking studies showing active site interactions of CouO and SgvM with L- and D-SAM as well as their respective substrates. Docking is based on the crystal structures of CouO with SAH (PDB: 5M58) and SgvM with SAH and Zn^2+^ (PDB: 8FTS). **A:** CouO in complex with 4,5,7-trihydroxy-3-phenylcoumarin and L-SAM, highlighting key residues involved in binding. **B:** CouO with the same substrate and D-SAM, showing altered cofactor interactions. **C:** SgvM bound to 4-methyl-2-oxovalerate, Zn^2+^, and L-SAM, illustrating active site coordination. **D:** SgvM with D-SAM and substrate, depicting changes in cofactor orientation.

Docking studies based on the SgvM crystal structure in complex with SAH and the tested substrate 4-methyl-2-oxovalerate (PDB: 8FTS)^[18]^ were also performed with L-SAM and D-SAM (Figure S17). Unlike CouO, which employs R116 to stabilise the SAM carboxylate, SgvM lacks a comparable positively charged residue. Instead, stabilisation of SAM is mediated mainly by interactions with the adenosyl unit and by Y150 interacting with the carboxyl group of the Met moiety. For SAM docked into the active site, the adenosyl units are superimposable. It seems that S141 can form an additional interaction with the carboxylate of the D-Met moiety which is distorted in comparison to docked L-SAM (Figure 3C and 3D). Our results indicate similar distortions for D-SAM to those observed in the docked CouO complex (Figure S17 C). The substrate is stabilised in the active site by coordination to a Zn^2+^-ion and by hydrogen bonding with the side chains of R132 and W104. As only the carboxyl and carbonyl groups function as anchor points, the remaining substrate atoms retain flexibility. Thus, the conformational adaptability of the substrate presumably enables a catalytically competent geometry relative to the cofactor, enabling D-SAM to participate in methyl transfer despite the less favourable orientation. In the active site of SgvM, the angle between the nucleophilic carbon of the substrate, the methyl group, and the sulfonium sulfur is approximately 174.2° for L-SAM (Figure S19 A), closely approaching the ideal linear transition state geometry (~180°) associated with S_N_2-type methyl transfer. For D-SAM, this angle deviates more substantially from linearity (144.9°, Figure S19 B), consistent with the reduced reactivity observed experimentally. When considering the angle between the methyl group and the molecular plane of the substrate, D-SAM (81.1°, Figure S19 D) is slightly closer to the ideal 90° than L-SAM (102.2°, Figure S19 C), although the overall flexibility of the substrate appears to be more important for D-SAM turnover than this angle alone.

Our experimental and docking-based results suggest that, when a strict binding mode is required for catalytic activity, as in CouO, the use of D-SAM as a methyl donor becomes more challenging. While SgvM tolerates both stereoisomers, CouO strictly selects L-SAM, indicating that D-SAM compatibility is not universal among *C*-MTs. Structural flexibility of substrate and active site appears crucial to accommodate the inverted geometry of D-SAM. These findings suggest that altered SAM stereochemistry may be useful as an additional design element in engineering MTs with modified cofactor selectivity.

### Radical SAM Enzymes as Unusual D-SAM Acceptors

Radical SAM enzymes constitute a large and diverse superfamily that utilise a [4Fe-4S] cluster to reductively cleave SAM, thereby generating a 5′-deoxyadenosyl radical which initiates a wide range of radical-mediated transformations. Their catalytic mechanism relies on the precise bidentate coordination of the amino acid moiety of SAM to the iron–sulfur cluster, which imposes stringent stereochemical requirements on the cofactor. Having observed D-SAM compatibility in other SAM-dependent enzymes, we next explored its conversion by radical SAM enzymes. To assess D-SAM compatibility across mechanistically diverse enzymes, we selected the tyrosine lyase HydG from *Thermoanaerobacter italicus*, the tryptophan 2-*C*-MT TsrM from *Streptomyces laurentii*, the cobalamin-dependent radical SAM MT GenD1 from *Micromonospora echinospora*, and the glutamine C-MT (QCMT) from *Methanoculleus thermophilus*. Both the carboxylate and amino group of SAM are coordinated by the [4Fe-4S] cluster. Hence, SAM stereochemistry is expected to be critical for catalysis, although productive turnover with D-SAM cannot be excluded if alternative binding modes are possible.

D-SAM was produced *in situ* by a MAT enzyme. MTAN was not included in the experiments to allow analysis of byproduct formation. TsrM was tested for methylation of tryptophan (L-Trp), GenD1 for methylation of gentamicin A, HydG for cleavage of tyrosine and QCMT for methylation of a synthetic 10-mer peptide (HFGGSQRAGV).

As expected, activity with L-SAM was observed for HydG. In the D-SAM sample, small amounts of both *p*-cresol and the byproduct 5′-deoxyadenosine (DOA) were initially detected. However, these peaks were also present in the control reaction without Met and could therefore be attributed to HydG-bound L-Met, likely remaining from activity on intracellular L-Tyr during enzyme production in *E. coli*. The experiment was therefore repeated with enzymatically produced and purified D-SAM to avoid potential L-SAM formation from L-Met catalysed by the MAT enzyme. Under these conditions, no formation of *p*-cresol or DOA was observed, confirming the interpretation of the previous experiment. The lack of activity with D-SAM aligns with structural data revealing stringent stereochemical requirements for SAM binding in this class of enzymes.^[21,22]^

TsrM operates by a distinct, cobalamin-dependent, non-radical mode of catalysis rather than by the conventional radical SAM mechanism: First, the methyl group is transferred to cobalamin (Cbl) to yield methyl cobalamin (MeCbl). MeCbl is then attacked by the deprotonated substrate L-Trp, resulting in formation of 2-methyltryptophan (2-MeTrp) and Cbl(I). It has been suggested that a second SAM molecule is essential for the reaction, with its carboxylate functioning as a catalytic base to deprotonate L-Trp.^[6]^ For this role, correct placement of the carboxylate moiety is essential. Here, the reaction was directly conducted with purified D-SAM or L-SAM. While the L-SAM positive control showed >99% conversion, neither 2-MeTrp nor the byproduct SAH was observed with purified D-SAM (Figure 4). This highlights the importance of SAM stereochemistry for the TsrM mechanism. The stereochemistry potentially influences the correct placement of the carboxylate to function as a catalytic base.

**Figure 4.**
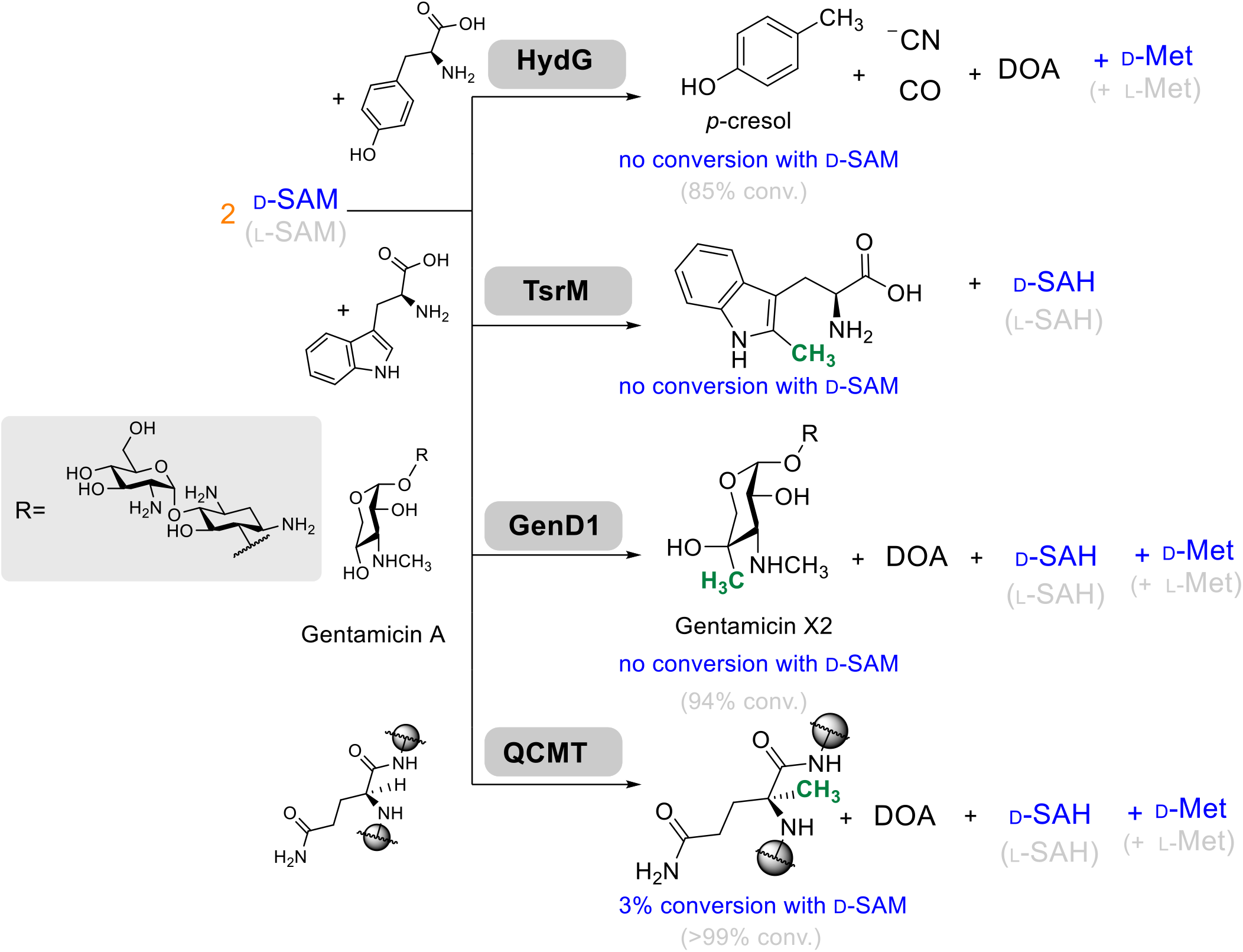
Reactivity of four different radical SAM enzymes with *in situ* SAM supply. Conversions with D-SAM are shown in blue and conversions with L-SAM are shown in grey. For QCMT and GenD1, two SAM molecules were required for one catalytic cycle (orange). QCMT was tested with a synthetic 10-mer peptide (HFGGS**Q**RAGV); for clarity, only the glutamine (Q) residue is shown. DOA = 5′-deoxyadenosine. Transferred methyl groups are shown in green in the methylated products.

GenD1 and QCMT, both radical SAM class B MTs, combine parts of the HydG and TsrM reactions utilising MeCbl to methylate unreactive carbon atoms *via* a substrate radical intermediate.^[23]^ Therefore, two SAM binding modes are essential for successful turnover: One for methylation of Cbl and one for coordination at the [4Fe-4S] cluster, allowing homolytic cleavage of SAM. Given that successful turnover requires two different productive SAM binding modes, no turnover with D-SAM was expected. No increase in product or byproduct formation was observed for GenD1 in the cascade reaction upon addition of D-Met compared to the no Met control. Curiously, QCMT catalysed methylation when D-Met was supplied, yielding the methylated product confirmed by LC-MS analysis (Figure S12 A). This result was further supported by experiments using purified D-SAM, in which peaks corresponding to the methylated peptide were detected only in the presence of both D-SAM and QCMT, whereas no such peaks were observed in the corresponding control reactions lacking SAM or QCMT (Figure S12 B). Although the conversion was low (3%), this stands in contrast with other radical SAM enzymes and suggests an unusual tolerance towards SAM diastereomers in QCMT. For turnover, SAM must bind in two different conformations: One for Cbl methylation and the other for SAM cleavage at the [4Fe-4S]-cluster. While alkylation of Cbl is a polar S_N_2 reaction and mechanistically less dependent on the precise conformation of SAM, the cleavage reaction is more challenging. The reduction potential of the reduced [4Fe-4S] cluster alone is not sufficient for reductive cleavage of SAM.^[24]^ The carboxyl and amino groups of SAM function as interaction partners to the [4Fe-4S]-cluster, likely tuning the reactivity in order to achieve the single electron transfer to SAM.^[21]^ It is therefore notable that this fine tuning can be achieved with a SAM methionyl analogue.

### Cyclases AZC Synthase and ACC Synthase

The SAM-dependent enzymes investigated so far accepted D-SAM either as a methyl donor or as the source of a SAM derived radical, without transferring the cofactor’s α-amino stereocentre to the product. Based on the broad acceptance of D-SAM across mechanistically diverse enzymes, we subsequently examined a SAM-dependent reaction in which the amino acid side chain rather than the methyl group is transferred to the product. In this reaction, the amino group attacks the Cγ atom of the Met-derived side chain (Figure 5). *Pa*AzeJ from *Pseudomonas aeruginosa* is a SAM-dependent enzyme that catalyses the intramolecular cyclisation of the amino acid side chain of L-SAM to yield L-azetidine-2-carboxylic acid (L-AZC) alongside MTA. Hence, when D-SAM is used, the enantiomeric D-azetidine-2-carboxylic acid (D-AZC) should be formed. Unlike in the earlier cases, the enzyme must not only bind the SAM diastereomer but also catalyse cyclisation of the Met-derived side chain, making this a particularly direct test of whether the stereochemistry of D-SAM is compatible with productive turnover. *Pa*AzeJ was applied in the established three-enzyme cascade employing *Uu*MAT, starting from either D- or L-Met. The two products were derivatised using Marfey’s reagent,^[25]^ and analysed against commercially available authentic standards (Figure 5). Derivatisation was performed in excess, yielding the expected diastereomeric products: the (L,L)-Marfey derivative from L-AZC formed from L-SAM and the (D,L)-Marfey derivative from D-AZC formed from D-SAM.

**Figure 5.**
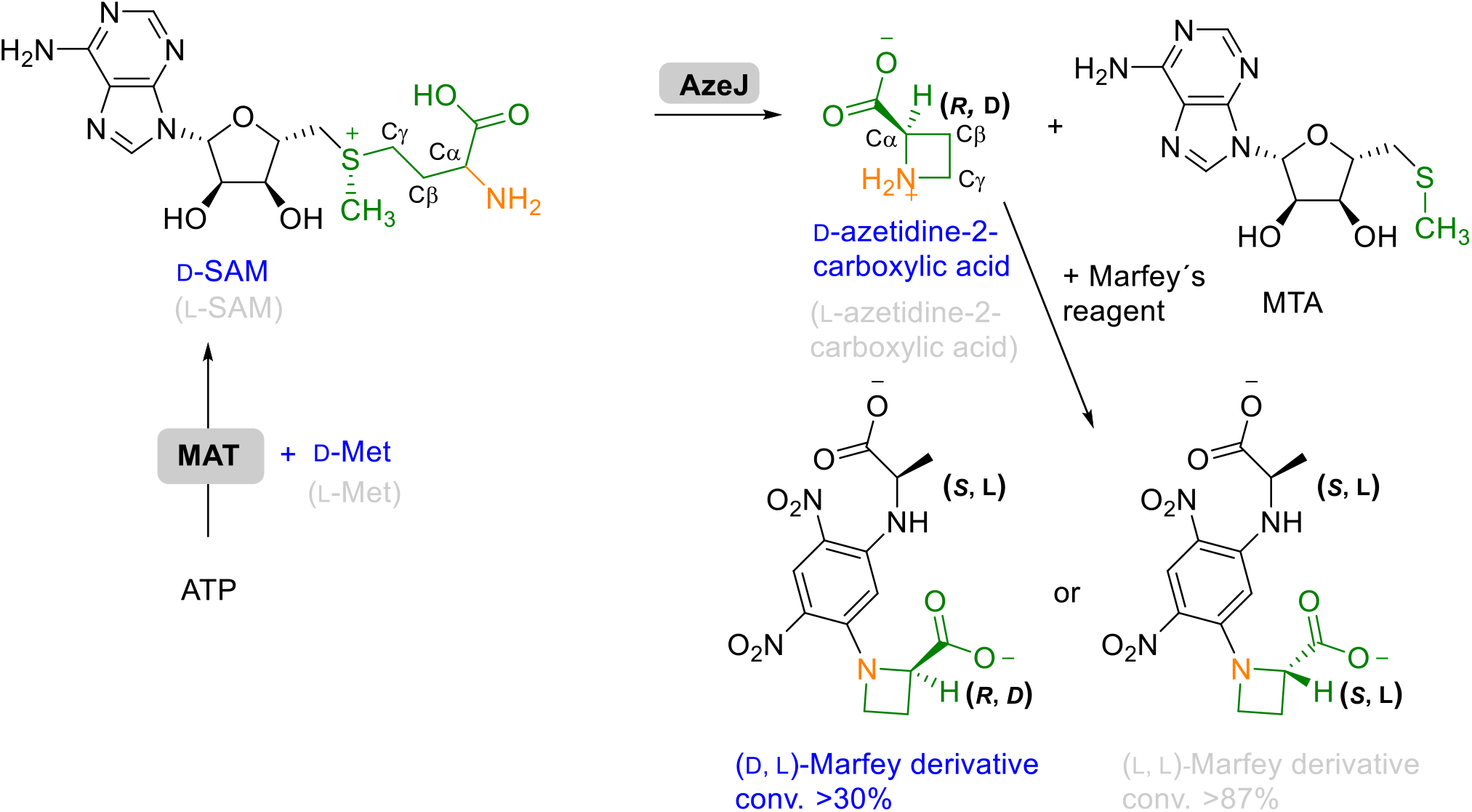
Conversion of D-SAM to D-azetidine-2-carboxylic acid and MTA. Adenosine-5′-triphosphate (ATP) and D-/L-met are converted to D- (blue) or L-SAM (grey) by methionine adenosyltransferase (MAT). The enzyme *Pa*AzeJ catalyses the cyclisation of SAM to azetidine-2-carboxylic acid and MTA. The resulting amino acid product is subsequently derivatised using Marfey’s reagent for stereochemical analysis. The Met moiety of SAM and the products and substituents derived from it are shown in green.

In the three-enzyme cascade, formation of the derivatised product starting from L-Met reached over 87%, while product formation starting from D-Met reached over 30%. As our primary objective here was to establish the qualitative feasibility of forming L- or D-AZC from L- or D-SAM, respectively, the reaction conditions were not further refined. While additional optimisation might increase the conversion of D-Met and hence D-SAM, our current conditions were sufficient to demonstrate the stereochemical outcome of the transformation.

These results confirm the formation of enantiomeric AZC from D- and L-SAM. Comparable D-SAM-dependent product formation was also observed with *Sac*AzeJ from *Saccharotrix sp*. NRRL B-16348, whose crystal structure was determined in this study (PDB: 29ST). The data suggest that conversion from D-SAM is lower than from L-SAM, although this comparison is limited by the influence of the subsequent derivatisation step and by the possibility that D-SAM formation in the cascade is less efficient than for L-SAM. To further investigate how D-SAM can act as a substrate, we performed docking studies to compare the binding modes of D-SAM and L-SAM within the active site of *Pa*AzeJ. The simulations were performed with the crystal structure of *Pa*AzeJ in complex with MTA and AZC (PDB: 8RYE)^[14]^. This structure is in good alignment to the SacAzeJ structure, overall sequence identity is 60% with an alignment r.m.s.d. of 0.87 Å (Figure S22). D-SAM can adopt a conformation in which its amino group is positioned on the opposite side of the binding pocket compared to L-SAM, resulting in a shorter distance to the γ-carbon atom (2.6 Å for D-SAM versus 2.9 Å for L-SAM), consistent with a geometry suitable for intramolecular cyclisation. Docking revealed multiple possible binding poses of D-SAM within the active site. The energetically preferred pose lacks the cation–π interaction, between F134 and the sulfonium, which has previously been proposed by Klaubert *et al*. to play a critical role in catalysis with L-SAM. Alternative poses that allow interaction with F134, exhibit suboptimal geometry for nucleophilic attack on the γ-carbon.^[14]^ Therefore, the preferred pose is likely to be catalytically relevant, suggesting that the absence of the interaction with F134 may reduce transition state stabilisation and thereby contribute to the lower conversion observed experimentally, while also indicating that catalysis with D-SAM may occur without assistance from F134 (Figure S20).

In contrast to AzeJ, 1-aminocyclopropane-1-carboxylic acid synthase (ACCS) from *Malus domestica* catalyses cyclisation of SAM into a three-membered ring amino acid using PLP. In this transformation, the α-amino group of SAM displaces the lysine-bound internal aldimine to form an external aldimine with PLP, thereby initiating cyclisation via PLP-mediated chemistry (Figure 6).^[26,27]^

**Figure 6.**
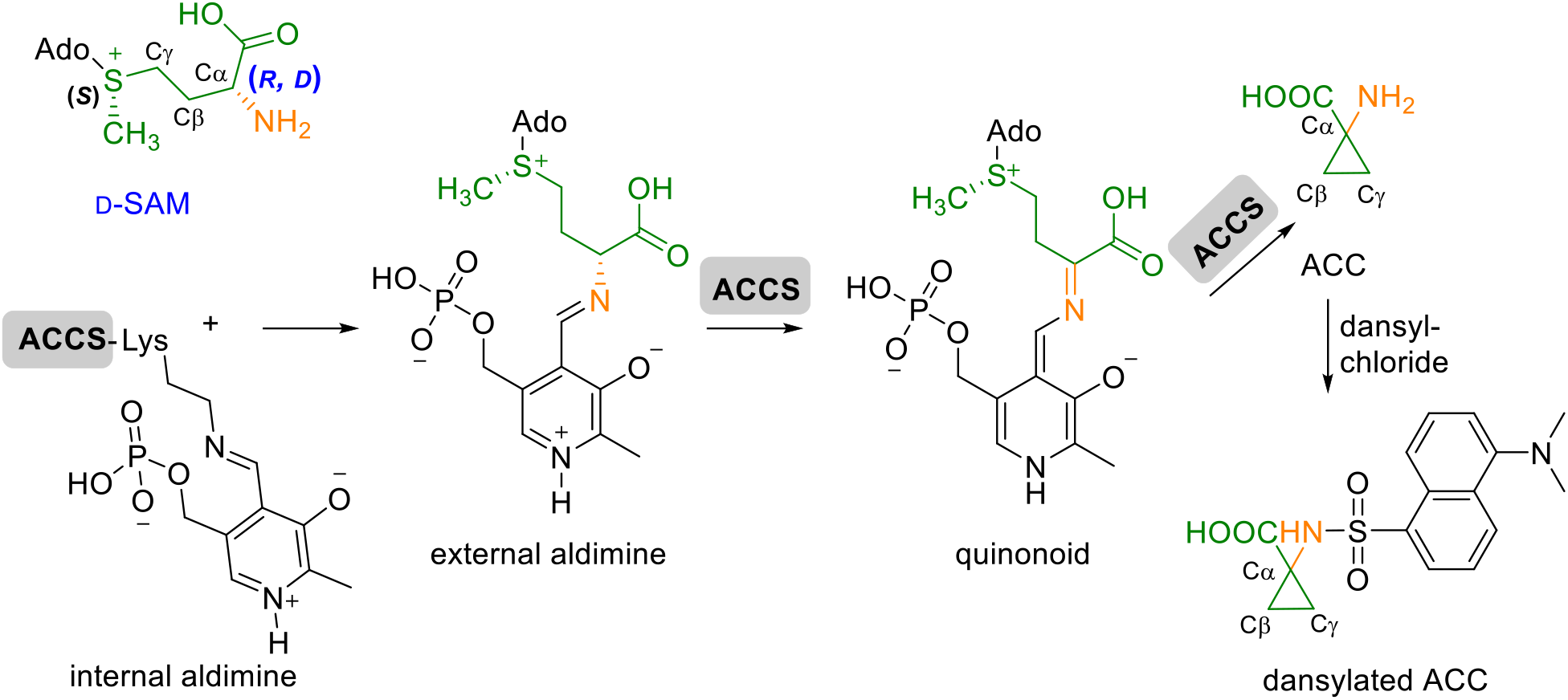
PLP-dependent conversion of L- or D-SAM to 1-aminocyclopropane-1-carboxylic acid (ACC) by ACC synthase (ACCS). The reaction proceeds *via* formation of an external aldimine from the internal aldimine, followed by quinoid intermediate formation and subsequent cyclisation to ACC. The resulting product was derivatised with dansyl chloride to yield dansylated ACC for detection.

The stereochemistry at the α-carbon of SAM is ultimately lost during the catalytic cycle, this occurs upon formation of the quinonoid intermediate. At this stage, the α-carbon becomes sp^2^-hybridised and planar. As a result, this intermediate should be identical regardless of whether it originates from D-SAM or L-SAM. The key question is whether the initial imine formation between PLP and the α-amino group of D-SAM can occur, given its distinct stereochemical arrangement. Using HPLC-MS/MS, product formation from D-SAM could indeed be detected, however, the signal corresponding to ACC is approximately fourfold lower compared to the reaction with L-SAM. Comparison with a dansylated reference compound of ACC confirmed the identity of the product: both the molecular ion at *m*/*z* 335.1 in positive ion mode and the characteristic fragmentation pattern matched those of the authentic standard. This observation is particularly surprising given that ACCS has been thought to require very strict geometric positioning of the α-amino group for catalysis, highlighting an unexpected flexibility in substrate stereochemistry.^[28]^ To further rationalise the detectable turnover of D-SAM by ACCS, docking studies with L- and D-SAM were performed based on the crystal structure of ACCS in complex with an amino oxy analogue (AMA) and PLP (PDB: 1M4N)^[29]^, together with the PLP-bound dimeric structure of ACCS (PDB: 1IAX)^[27]^. For L-SAM, the docked conformation shows a favourable orientation of the Cα-H-bond towards K273 (2.7 Å), consistent with direct deprotonation by this residue. By comparison, D-SAM adopts a conformation in which the Cα-H bond points away from K273, indicating that the canonical deprotonation mode is less favourable. At the same time, the docked D-SAM conformation places the Cα-H bond close to the SAM carboxylate (2.6 Å), while the carboxyl group remains close to K273 (2.8 Å). Although this does not yet establish the mechanism, it suggests that alternative proton abstraction pathways could be possible. Moreover, formation of the PLP-bound imine is expected to increase the acidity of the α-carbon and may therefore help to enable proton abstraction even in the less favourable D-SAM binding mode (Figure S21). We are currently investigating this further, as the ACCS result may also have implications for other PLP-dependent enzymes acting on SAM-derived amino acid substrates.

## Conclusion

Our results demonstrate that the tested model *N*- and *O*-MTs, as well as one of the examined *C*-MTs, can utilise D-SAM, albeit typically with lower conversions compared to canonical L-SAM. Especially for *C*-MTs, the angle between the transferable methyl group of SAM and the nucleophilic substrate carbon, as well as substrate flexibility within the active site, appear to be crucial factors for productive turnover with D-SAM. In CouO, the R116 residue likely contributes to stereoselectivity by stabilising the carboxylate group of L-SAM while disfavouring productive binding of D-SAM. Protein engineering approaches targeting such residues may help improve activity with D-SAM. Our molecular docking studies of *C*-MTs thus provide mechanistic insights into how altered SAM stereochemistry can be accommodated or rejected in different active-site environments.

Beyond classical MTs, our study also explored the use of D-SAM in radical SAM enzymes. Most enzymes, including HydG, TsrM, and GenD1, showed no detectable turnover with D-SAM. For HydG, this is consistent with the stringent stereochemical requirements for SAM binding to the [4Fe-4S] cluster, whereas in TsrM and GenD1 additional constraints arise from their more complex cobalamin-dependent mechanisms. In contrast, QCMT exhibited low but measurable activity. This unexpected conversion suggests that QCMT can accommodate D-SAM despite its dual requirement for cobalamin methylation and radical SAM chemistry, indicating some degree of cofactor tolerance or active site flexibility.

For the PLP-dependent ACCS, conversion of D-SAM was not expected, as the α-amino group is directly involved in the catalytic mechanism. Nevertheless, D-SAM was accepted as a substrate. Similarly, the cyclase AzeJ catalyses the formation of AZC with a new stereogenic centre from D-SAM. Again, the conversion is lower than observed for L-SAM. Importantly, when the Met-derived side chain of SAM is directly incorporated into the product, as in the AzeJ-catalysed reaction, the stereogenic centre of D-SAM can be transferred. This may provide opportunities to expand product diversity and could be relevant for the development of partially orthogonal enzymatic systems. Together with our results on classical and radical SAM enzymes, these insights broaden the known scope of enzyme compatibility with D-SAM and underscore nature’s versatility in cofactor recognition. An overview of SAM-dependent enzymes investigated with D-SAM, including the enzymes examined in this study, is given in Table S10.

Building on earlier studies, our findings extend the current understanding of D-SAM utilisation and identify first starting points for engineering enzyme variants with shifted cofactor preference between L-SAM and D-SAM. In the longer term, this could contribute to the development of biocatalytic pathways with tailored cofactor selectivity, and complement recent metabolic engineering efforts to expand SAM biosynthesis *in vivo*.^[30]^ Moreover, recent progress in enzymatic *in situ* synthesis of L-Met analogues to generate SAM analogues suggest promising opportunities to develop similar systems for D-Met analogues, enabling efficient production and utilisation of D-SAM analogues. In a biocatalytic context, acceptance of D-Met or racemic Met mixtures could simplify substrate supply and provide access to alternative SAM diastereomers without prior enantiomeric separation. Given that D-Met is known to occur in bacterial systems,^[31]^ it will be interesting to examine whether D-SAM may also arise under specific physiological conditions, which would further expand the known scope of cofactor diversity, although its natural occurrence has not yet been established.

## Acknowledgements

We thank the Deutsche Forschungsgemeinschaft (DFG, FOR 5596, project 510974120) for financial support. Z.Z. thanks the Chinese Scholarship Council (CSC) for a Ph.D. fellowship. We thank Prof. Squire J. Booker (PennState, USA) for providing pBAD42_btuCEDFB, Prof. Peter Roach (Southampton, UK) for providing pRD003_hydg, and Prof. Andrea Rentmeister (LMU München, Germany) for providing pET-28a_couo. We acknowledge DESY (Hamburg, Germany), a member of the Helmholtz Association HGF, for the provision of experimental facilities. Parts of this research were carried out at beamline P11 at the PETRA III storage ring and we would like to thank the beamline staff for assistance during data collection. Beamtime was allocated for proposal Xh-20010031.

## Footnotes

1 For clarity and practicality, we refer to the diastereomeric forms as D-SAM and L-SAM throughout this study. Despite the D- and L-notation, these are not enantiomers, as only the stereocentre at the α-carbon is inverted, while the configurations at the ribose moiety and sulfonium centre remain unchanged.

## References

[1] P. Germer, J. N. Andexer, M. Müller, Synthesis 2022, 54, 4401; b) G. L. Cantoni, Annu. Rev. Biochem. 1975, 44, 435; c) M. K. F. Mohr, A. Satanowski, S. N. Lindner, T. J. Erb, J. N. Andexer, Microb. Cell Fact. 2025, 24, 55.

[2] H. Ü. Kaniskan, M. L. Martini, J. Jin, Chem. Rev. 2018, 118, 989; b) S. Wang, S. O. Klein, S. Urban, M. Staudt, N. P. F. Barthes, D. Willmann, J. Bacher, M. Sum, H. Bauer, L. Peng et al., Nat. Commun. 2024, 15, 43.

[3] Q. Sun, M. Huang, Y. Wei, Pharm. Sin. B 2021, 11, 632.

[4] M. Fontecave, M. Atta, E. Mulliez, Trends Biochem. Sci. 2004, 29, 243.

[5] a) J. B. Broderick, W. E. Broderick, B. M. Hoffman, FEBS Lett. 2023, 597, 92; b) P. A. Frey, A. D. Hegeman, F. J. Ruzicka, Crit. Rev. Biochem. Mol. Biol. 2008, 43, 63.

[6] H. L. Knox, P. Y.-T. Chen, A. J. Blaszczyk, A. Mukherjee, T. L. Grove, E. L. Schwalm, B. Wang, C. L. Drennan, S. J. Booker, Nat. Chem. Biol. 2021, 17, 485.

[7] a) Z. Hong, A. Bolard, C. Giraud, S. Prévost, G. Genta-Jouve, C. Deregnaucourt, S. Häussler, K. Jeannot, Y. Li, Angew. Chem. Int. Ed. 2019, 58, 3178; b) L. Barra, T. Awakawa, I. Abe, JACS Au 2022, 2, 1950.

[8] a) G. L. Cantoni, J. Biol. Chem. 1953, 204, 403; b) G. L. Cantoni, J. Am. Chem. Soc. 1952, 74, 2942.

[9] a) F. Schlenk, C. H. Hannum, A. J. Ferro, Arch. Biochem. Biophys. 1978, 187, 191; b) F. Schlenk, R. E. DePalma, J. Biol. Chem. 1957, 229, 1037.

[10] J. L. Galman, F. Parmeggiani, L. Seibt, W. R. Birmingham, N. J. Turner, Angew. Chem. Int. Ed. 2022, 61, e202112855.

[11] a) R. T. Borchardt, Y. Shiong, J. A. Huber, A. F. Wycpalek, J. Med. Chem. 1976, 19, 1104; b) K. D. Nakamura, F. Schlenk, Arch. Biochem. Biophys. 1976, 177, 170.

[12] J. Breiltgens, Z. Zou, J. N. Andexer, M. Müller, ChemBioChem 2026, DOI: 10.1002/cbic.70294.

[13] E. Abdelraheem, B. Thair, R. F. Varela, E. Jockmann, D. Popadić, H. C. Hailes, J. M. Ward, A. M. Iribarren, E. S. Lewkowicz, J. N. Andexer et al., ChemBioChem 2022, 23, e202200212.

[14] T. J. Klaubert, J. Gellner, C. Bernard, J. Effert, C. Lombard, V. R. I. Kaila, H. B. Bode, Y. Li, M. Groll, Nat. Commun. 2025, 16, 1348.

[15] E. Jockmann, F. Subrizi, M. K. F. Mohr, E. M. Carter, P. M. Hebecker, D. Popadić, H. C. Hailes, J. N. Andexer, ChemCatChem 2023, 15, e202300930.

[16] E. Jockmann, H. Girame, W. Steinchen, K. Kind, G. Bange, K. Tittmann, M. Müller, F. Feixas, M. Garcia-Borràs, J. N. Andexer, ACS Catal. 2025, 15, 6410.

[17] a) H. Lin, Bioorg. Chem. 2011, 39, 161; b) X.-W. Zou, Y.-C. Liu, N.-S. Hsu, C.-J. Huang, S.-Y. Lyu, H.-C. Chan, C.-Y. Chang, H.-w. Yeh, K.-H. Lin, C.-J. Wu et al., Acta Crystallogr. D 2014, 70, 1549.

[18] S. Ju, K. P. Kuzelka, R. Guo, B. Krohn-Hansen, J. Wu, S. K. Nair, Y. Yang, Nat. Commun. 2023, 14, 5704.

[19] a) C. Sommer-Kamann, J. Breiltgens, Z. Zou, S. Gerhardt, R. Saleem-Batcha, F. Kemper, O. Einsle, J. N. Andexer, M. Müller, ChemBioChem 2024, 25, e202400258; b) C. Sommer-Kamann, A. Fries, S. Mordhorst, J. N. Andexer, M. Müller, Angew. Chem. Int. Ed. 2017, 56, 4033.

[20] T. Pavkov-Keller, K. Steiner, M. Faber, M. Tengg, H. Schwab, M. Gruber-Khadjawi, K. Gruber, PloS one 2017, 12, e0171056.

[21] J. B. Broderick, B. R. Duffus, K. S. Duschene, E. M. Shepard, Chem. Rev. 2014, 114, 4229.

[22] R. D. Britt, L. Tao, G. Rao, N. Chen, L.-P. Wang, ACS Bio Med Chem Au 2022, 2, 11.

[23] a) J. Gagsteiger, S. Jahn, L. Heidinger, L. Gericke, J. N. Andexer, T. Friedrich, C. Loenarz, G. Layer, Angew. Chem. Int. Ed. 2022, 61, e202204198; b) C. Huang, F. Huang, E. Moison, J. Guo, X. Jian, X. Duan, Z. Deng, P. F. Leadlay, Y. Sun, Chem. Biol. 2015, 22, 251; c) A. Benjdia, O. Berteau, Curr. Opin. Struct. Biol. 2023, 83, 102725.

[24] M. W. Ruszczycky, H.-w. Liu, Biochemistry 2024, 63, 3161.

[25] P. Marfey, Carlsberg Res. Commun. 1984, 49, 591.

[26] a) G. Capitani, D. L. McCarthy, H. Gut, M. G. Grütter, J. F. Kirsch, J. Biol. Chem. 2002, 277, 49735; b) Y. Li, L. Feng, J. F. Kirsch, Biochemistry 1997, 36, 15477.

[27] G. Capitani, E. Hohenester, L. Feng, P. Storici, J. F. Kirsch, J. N. Jansonius, J. Mol. Biol. 1999, 294, 745.

[28] S. Khani-Oskouee, J. P. Jones, R. W. Woodard, Biochem. Biophys. Res. Commun. 1984, 121, 181.

[29] G. Capitani, A. C. Eliot, H. Gut, R. M. Khomutov, J. F. Kirsch, M. G. Grütter, Biochim. Biophys. Acta 2003, 1647, 55.

[30] a) M. K. F. Mohr, R. Saleem-Batcha, N. V. Cornelissen, J. N. Andexer, Chem. Eur. J. 2023, 29, e202301503; b) M. K. F. Mohr, P. Benčić, J. N. Andexer, Angew. Chem. Int. Ed. 2025, 64, e202414598.

[31] A. Aliashkevich, L. Alvarez, F. Cava, Front. Microbiol. 2018, 9, 683.

