## Supplementary material for "S-Adenosyl-D-methionine as a Non-Physiological Substrate for a Wide Range of SAM-Dependent Enzymes": SI

### 1. General and Experimental

#### 1.1 Chemicals and Solvents

Chemicals and solvents were purchased from Sigma-Aldrich, Carl Roth or Thermo Scientific unless otherwise specified. Solvents were obtained either anhydrous or in HPLC grade.

#### 1.2 Plasmids, Protein Overproduction, Protein Purification and Reconstitution of Radical SAM Enzymes

The plasmids *Tk*MAT, *Rg*ANMT, *Pp*CaOMT, CouO, SgvM, *Ec*MTAN, AzeJ, ACCS, HydG, *Uu*MAT, QCMT, as well as the corresponding protocols for protein overproduction and purification, have been described in previous publications.[1,2,3] Additional procedures relevant to the present study are described below. Radical SAM enzymes were handled under anaerobic conditions and reconstituted after purification.

*Protein Overproduction of TsrM and GenD1*

The plasmid pRD003_hydg, harbouring hydg from Thermoanaerobacter italicus together with the isc operon from E. coli, served as the backbone for construction of pRD003_genD1 and pRD003_tsrM.[3] The pRD003 vector was linearised by restriction digestion using NcoI and XhoI. The respective genes were amplified by PCR from synthetic DNA using primers introducing a 5′ NcoI restriction site (CCATGG) and a 3′ XhoI restriction site (CTCGAG). For production of TsrM and GenD1, cultures were grown in 1.3 L LB medium supplemented with the appropriate antibiotics in non-baffled 2 L or 3 L flasks, inoculated with 13 mL seed culture, and incubated at 37 °C and 120 rpm. At inoculation, cultures were supplemented with 2 mM ammonium ferric citrate and 2 mM L-cysteine. For TsrM, hydroxocobalamin (1.3 µM) was added additionally. Expression of pRD003-based constructs (pRD003_*genD1 and* pRD003_*tsrM*) was induced with 0.2% (w/v) D-arabinose. Cultures were then incubated at 20 °C and 120 rpm. Cell pellets were stored at -80 °C until use. For production of TsrM, the pRD003_tsrm vector was cotransformed with pBAD42_*btu*CEDFB.[4]

*Purification of TsrM and GenD1*

Purification of TsrM and GenD1 was performed under strictly anaerobic conditions and differed from the general protein purification procedure in the following aspects. All steps were carried out in an anaerobic chamber (0 ppm $O_2$, ≥2.0% $H_2$) or in sealed containers maintaining the chamber atmosphere. Materials were transferred into the chamber at least 24 h before use, and all buffers and solutions were rendered anaerobic prior to use. Cell pellets were thawed and resuspended in lysis buffer (3 mL $g^{-1}$ wet cell mass) and disrupted by bead beating using 2 cycles at 6000 rpm for 20 s with 5 min cooling on ice between cycles. The crude lysate was clarified by centrifugation at 50,873 × g for 45 min at 4 °C. Proteins were purified from the cleared lysate by Ni-NTA chromatography as described for the general purification procedure, except that buffer A was used for the first wash, followed by buffer B containing 20 mM imidazole, and, for GenD1, an additional wash with buffer B containing 50 mM imidazole (Table 1). Desalting was performed using a PD-10 column with buffer A. Protein concentrations were determined by NanoDrop UV/Vis spectroscopy, and purified proteins were stored under anaerobic atmosphere at 4 °C prior to reconstitution.

Table S1: Buffers used for purification of radical SAM enzymes.
*Buffer B was diluted with buffer A to reach different imidazole concentrations

| Buffer | Component | Concentration |
|---|---|---|
| Lysis buffer, buffer A, pH = 7.9 | Tris-HCl | 20 mM |
| | NaCl | 500 mM |
| | Glycerol | 10% (V/V) |
| Buffer B* pH = 7.9 | Tris-HCl | 20 mM |
| | NaCl | 500 mM |
| | Imidazole | 1 M |
| | Glycerol | 10% (V/V) |
| Storage buffer pH = 7.9 | Tris-HCl | 20 mM |
| | NaCl | 250 mM |
| | Glycerol | 25% (V/V) |

*Reconstitution of radical SAM enzymes*

Reconstitution of [4Fe-4S] clusters was performed for all radical SAM enzymes on the day after purification under anaerobic conditions at room temperature. DTT was added to a final concentration of 5 mM, and the solution was incubated for 30 min. Ammonium iron(II) sulfate (Mohr's salt) was then added slowly in a 5-fold molar excess relative to the number of [4Fe-4S] clusters per protein, followed by incubation for 30 min. Subsequently, lithium sulfide was added in an amount equimolar to Mohr's salt, and the mixture was incubated for another 30 min. Formed FeS colloids were removed by centrifugation (4000 rpm, 30 min), and the protein solution was desalted by size-exclusion chromatography using a PD-10 column. Reconstituted proteins were concentrated by spin filtration (30 kDa MWCO), aliquoted (50–100 µL), snap-frozen in liquid nitrogen, and stored at −80 °C until use.

##### 1.3 Enzymatic Preparation of D-SAM and L-SAM

Preparative synthesis of L-SAM and D-SAM was carried out using *Tk*MAT in a total volume of 3 mL containing 10 µM *Tk*MAT, 50 mM Tris-HCl (pH 7.5), 20 mM $MgCl_2$, 50 mM KCl, 6 mM ATP, and 6 mM L-Met or D-Met, respectively. Reactions were incubated at 37°C and 350 rpm for 4 h (L-Met) or 6 h (D-Met). Reactions were quenched by addition to HCl to a final concentration of 300 mM and centrifuged for 15 min at 12,700 rpm.

The supernatant was filtered (0.2 µm) and diluted to 10 mL prior to purification by strong cation exchange (SCX) chromatography on a HiTrap SP FF 1mL column using 50 mM ammonium formate (pH 4.0) as mobile phase A and 700 mM ammonium formate as mobile phase B at a flow rate of 1 mL $min^{-1}$. SAM-containing fractions were combined, frozen in liquid nitrogen and lyophilised.

For desalting, the lyophilised material was dissolved in 110 µL 100 mM HCl and purified by reversed-phase HPLC on an ISERA ISAspher 100-5 C18 AQ column (250 x 8 mm) using 0.1% formic acid in water as mobile phase A and MeCN as mobile phase B at 1 mL $min^{-1}$. SAM-containing fractions were combined, frozen in liquid nitrogen, and lyophilised. A schematic overview of the purification procedure and representative chromatograms of purified L-SAM and D-SAM are shown in Figure S1. The purified material was dissolved in 0.1 M HCl, quantified by HPLC method B (see section 1.4.3, Table S4), and stored at -80 °C until use.

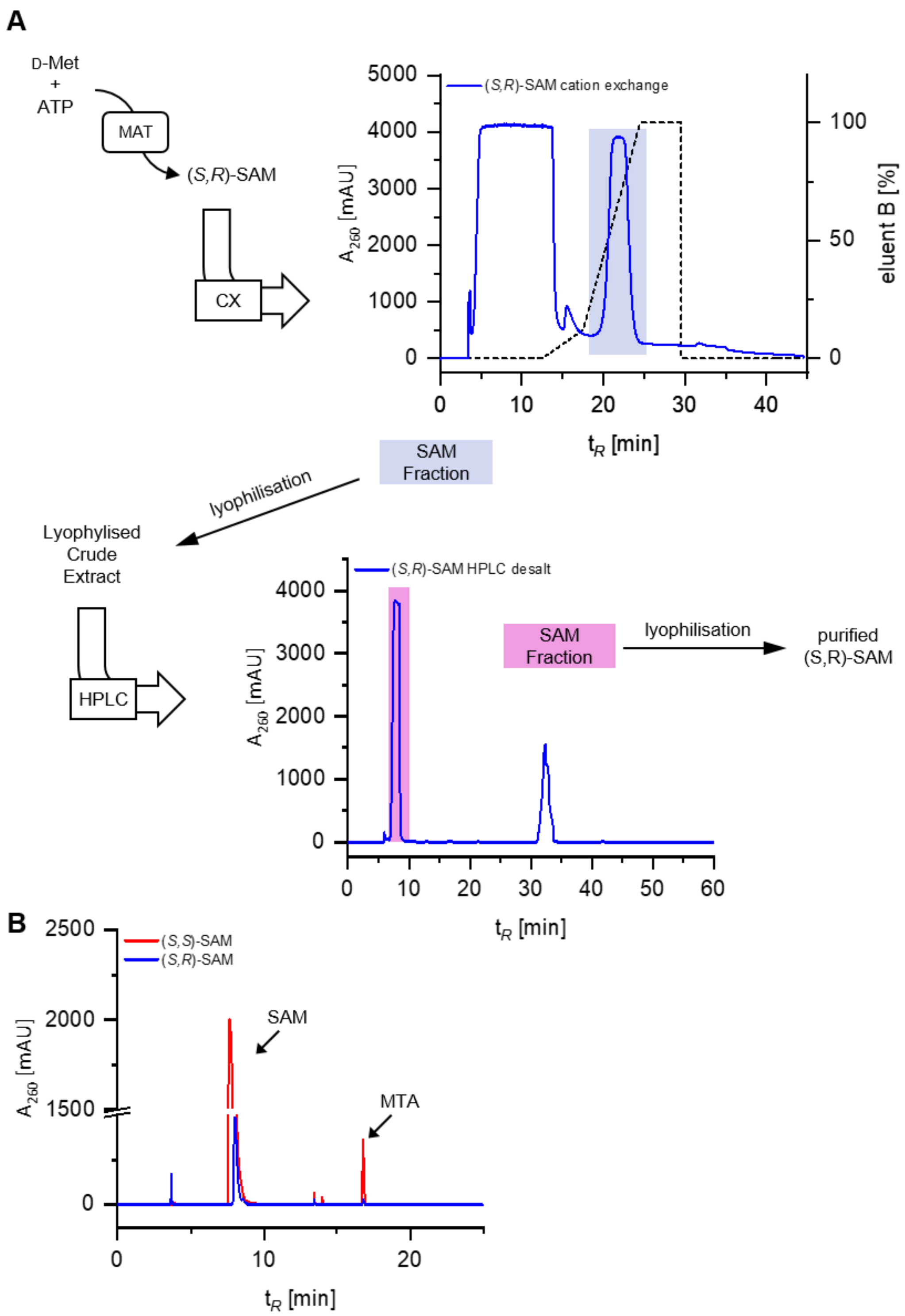

Figure S1: Purification of L-SAM [(*S*,S)-SAM] and D-SAM [(*S*,*R*)-SAM]. (A) Schematic overview of the purification procedure. The MAT reaction mixture was first subjected to cation exchange (CX) chromatography. Fractions highlighted in blue were collected and lyophilised. The resulting crude material was then purified by RP-HPLC, and the peak highlighted in pink was collected and lyophilised to yield purified SAM. For preparation of L-SAM, L-methionine was used, whereas D-methionine was used for preparation of D-SAM. (B) HPLC chromatograms of purified L-SAM (red) and D-SAM (blue). Minor amounts of the SAM degradation product methylthioadenosine (MTA) are present in both samples, consistent with the intrinsic instability of SAM.

##### 1.4 Enzyme Assays and Analytical Methods (with Supporting Tables and Figures)

Enzyme activity was assayed using the physiological cofactor/substrate L-SAM to confirm catalytic activity under the applied conditions. In all cases, appropriate negative control reactions were performed. For comparison, analogous assays were then carried out using D-SAM, either generated in situ from D-Met and ATP or added directly after enzymatic preparation and purification, as specified for the respective enzyme system. For SgvM, AzeJ, and ACCS, analytes were derivatised prior to analysis. Where conversion values are given, they were calculated from chromatographic peak areas (AUC) according to

$$Conversion\ [\%] = \frac{AUC(Product)}{\mathrm{AUC}(Substrate) + AUC\ (Product)}\ x\ 100$$

Reactions in which no residual substrate peak was detectable were interpreted as showing >99% conversion.

###### *1.4.1 Assay Conditions for Methyltransferases RgANMT, PpCaOMT and CouO and HPLC Analysis*

Assays were conducted in 1.5 mL reaction tubes at 37 °C with shaking at 350 rpm using a Thermomixer Comfort (Eppendorf SE). Each reaction contained 50 mM TRIS buffer (pH 7.5), 20 mM $MgCl_2$ and 50 mM KCl. The enzymes and substrates were added at the following concentrations: methionine adenosyltransferase (*Tk*MAT) at 17 µM, methyltransferase (MT) at 3 µM, methylthioadenosine nucleosidase (*Ec*MTAN) at 1 µM, ATP at 1 mM, methionine at 1 mM, and substrate (2-amino-4-nitrophenol for *Rg*ANMT and *Pp*CaOMT; 4,5,7-trihydroxy-3-phenylcoumarin for CouO) at 0.5 mM. The final reaction volume was adjusted with deionised water.

HPLC analysis was performed using an Agilent 1260 Infinity II system (Agilent Technologies Inc.) equipped with an diode array detector (DAD, Agilent Technologies Inc.). Separation was carried out on an ISAspher 100-3 C18 column (ISERA) at a temperature of 25 °C. The mobile phase consisted of solvent A (0.1% v/v formic acid in water) and solvent B (MeCN). The injection volume was 5 µL. The gradient was set up as follows at a flow rate of 1.0 mL/min (25 °C): 0.00–4.00 min, 3% B; 4.00–7.00 min, 3% to 70% B; 7.00–7.10 min, 70% to 100% B; 7.10–10.00 min, 100% B; 10.00–11.00 min, 100% to 3% B; 11.00–14.00 min, 3% B.

*Supporting Tables and Figures (RgANMT, PpCaOMT and CouO)*

In Table S2, the substrates (of *Rg*ANMT, *Pp*CaOMT and CouO) and their corresponding methylated products are presented along with their respective retention times.

Table S2: Retention times of substrates and methylated products in *Rg*ANMT, *Pp*CaOMT and CouO assays.

| Compound | Retention time (min) |
|---|---|
| 2-Amino-4-nitrophenol (2,4-ANP) | 7.8 |
| *N*-methylated product | 8.6 |
| *O*-methylated product | 8.7 |
| 4,5,7-Trihydroxy-3-phenylcoumarin | 8.5 |
| *C*-methylated product | 8.7 |

The assay conditions for *Rg*ANMT, *Pp*CaOMT, and CouO are described above. Representative HPLC chromatograms for the assays of *Rg*ANMT, *Pp*CaOMT, and CouO with L- and D-methionine are shown in Figures S2 to Figure S3. Time course analyses depicting substrate conversion and enzyme activity are presented in Figures S4, S5 and Figure S6.

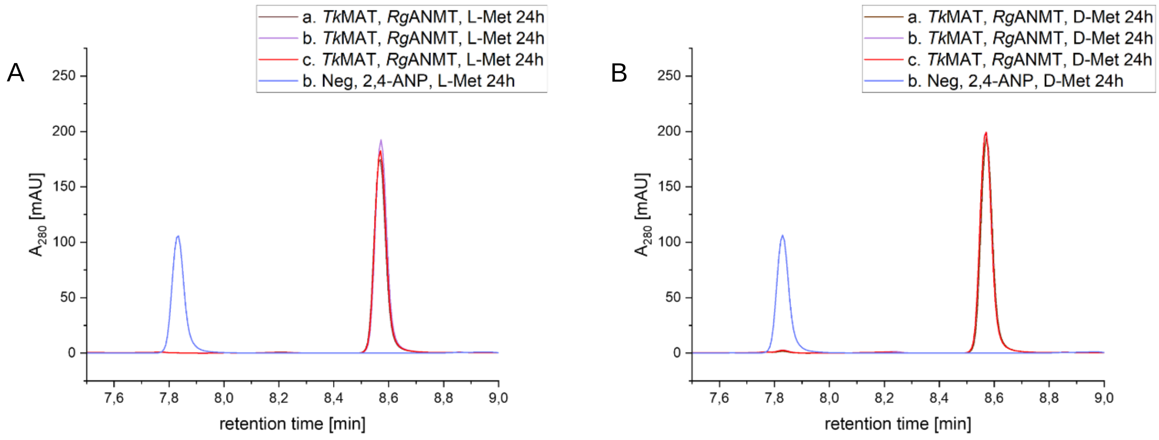

Figure S2: Chromatograms of *Rg*ANMT, *Tk*MAT and *Ec*MTAN. A**:** *Tk*MAT, *Rg*ANMT, L-Met triplicates after 24 h in brown, purple and red. Negative control (without enzymes) after 24 h in blue. B: *Tk*MAT, *Rg*ANMT, D-Met triplicates after 24 h in brown, purple and red. Negative control (without enzymes) after 24 h in blue.

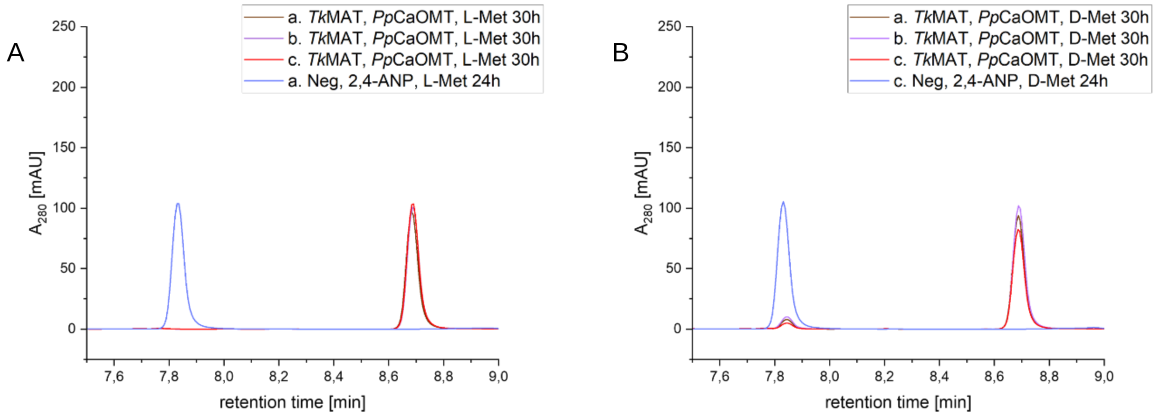

Figure S3: Chromatograms of *Pp*CaOMT, *Tk*MAT and *Ec*MTAN. A: *Tk*MAT, *Pp*CaOMT, L-Met triplicates after 30 h in brown, purple and red. Negative control (without enzymes) after 24 h in blue. B: *Tk*MAT, *Pp*CaOMT, D-Met triplicates after 30 h in brown, purple and red. Negative control (without enzymes) after 24 h in blue.

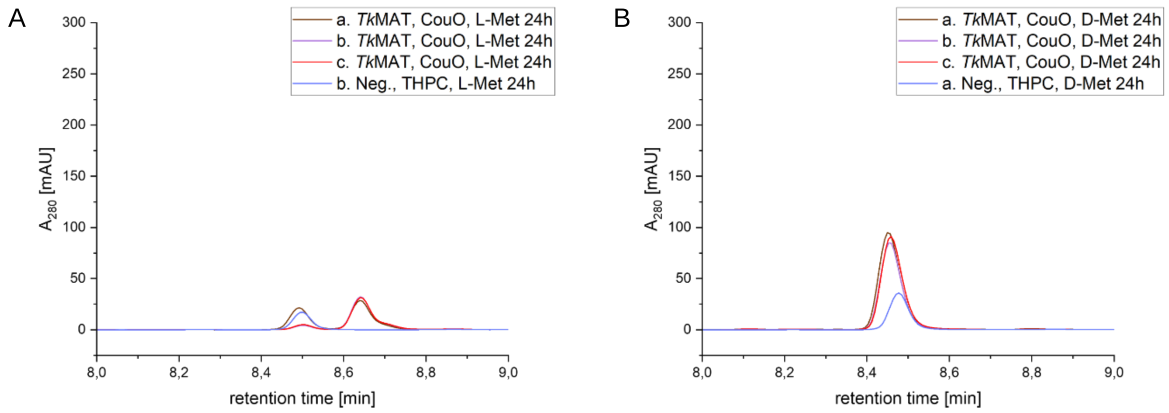

Figure S4: Chromatograms of CouO, *Tk*MAT and *Ec*MTAN. A: *Tk*MAT, CouO, L-Met triplicates after 24 h in brown, purple and red. Negative control (without enzymes) after 24 h in blue. B: *Tk*MAT, CouO, D-Met triplicates after 24 h in brown, purple and red. Negative control (without enzymes) after 24 h in blue.

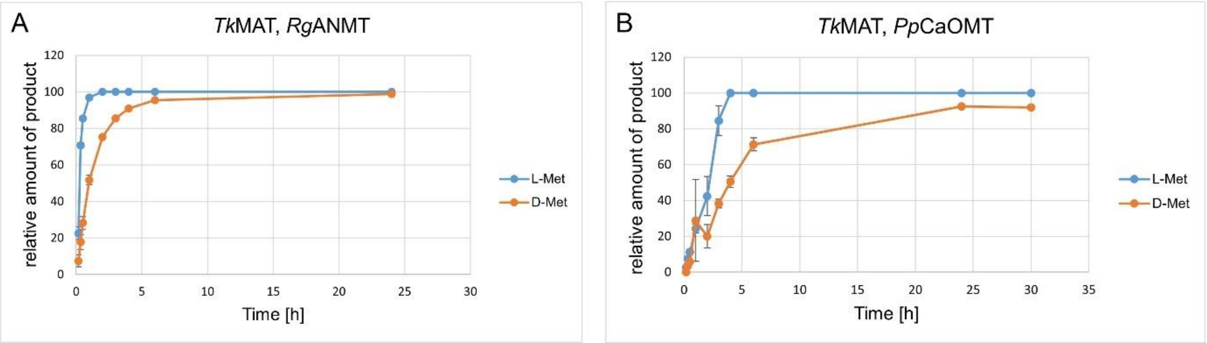

Figure S5: Time courses for *Rg*ANMT, *Pp*CaOMT and CouO. Time course of *Rg*ANMT (A) and *Pp*CaOMT (B) in three-enzyme cascade with *Tk*MAT. Error bars indicate standard deviation between triplicates. A: Blue: *Tk*MAT, *Rg*ANMT, L-Met. Orange: *Tk*MAT, *Rg*ANMT, D-Met. B: Blue: *Tk*MAT, *Pp*CaOMT, L-Met. Orange: *Tk*MAT, *Pp*CaOMT, D-Met.

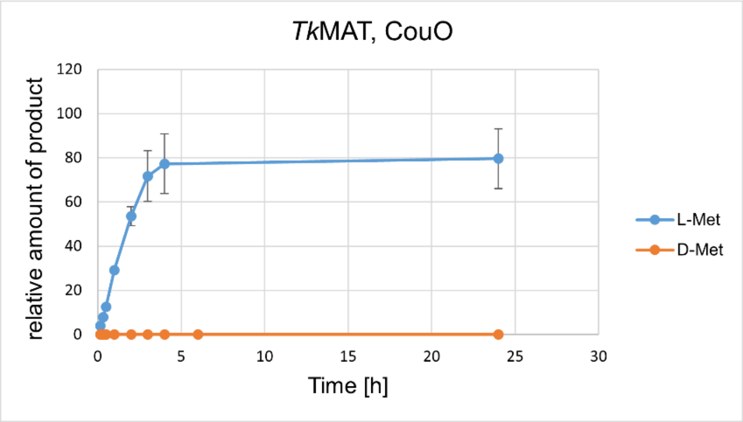

Figure S6: Time course of CouO in three-enzyme cascade with *Tk*MAT. Error bars indicate standard deviation between triplicates. Blue: *Tk*MAT, CouO, L-Met. Orange: *Tk*MAT, CouO, D-Met.

##### 1.4.2 *Assay Conditions for C-Methyltransferase SgvM and HPLC Analysis*

The enzyme assays with keto acids and SgvM were conducted according to the following protocol: 50 mM Tris-HCl pH 8.0, 50 mM $MgCl_2$, 20 mM KCl, substrate (1 mM), ATP (1.5 mM), L-methionine/D-methionine (1.5 mM), SgvM (0.8 mg·mL$^{-1}$), *Tk*MAT (0.5 mg·mL$^{-1}$), *Ec*MTAN (0.01 mg·mL$^{-1}$). The activity assays were incubated at 25 °C, 300 rpm for 20 h. For 4-methyl-2-oxovaleric acid, a time course analysis was performed with D- and L-methionine over 33 hours.

Detection after derivatisation: A 500 µL assay was incubated as described above. After the addition of 250 µL freshly prepared *O*-phenylenediamine (OPDA) in HCl (10 mg · mL$^{-1}$ in 1 M HCl), the mixture was incubated at 35 °C, 500 rpm for 45 min.[5] Substrates or products, if present, were extracted three times with 250 µL ethyl acetate and centrifuged for 2 min at 8000 rpm. The extracted phase was collected in a separate tube and subsequently concentrated in the concentrator (Concentrator 5301, Eppendorf, Hamburg, DE) under vacuum. Samples were then redissolved in methanol for HPLC analysis conducted by reversed-phase HPLC on an Agilent Technologies 1260 infinity II HPLC system with an Agilent Eclipse XDB-C18 column (250 mm × 4.4 mm, 5 µm, Agilent, Germany). Mobile phase A was $H_2O$ (0.1% formic acid), and mobile phase B was MeCN. The gradient was set up as follows at a flow rate of 0.5 mL/min (23 °C): 2-30 min 10% B to 80% B, 31-33 min 80% B, 34-40 min 80% to 10% B, 40-42 min 10% B. Peaks were detected using a DAD at 230-280 nm.

*Supporting Tables and Figures (SgvM)*

The corresponding retention times for substrates and methylated products are summarised in Table S3. Product formation in SgvM assays with 4-methyl-2-oxovaleric acid and 2-oxovaleric acid was confirmed by HPLC analysis (Figure S7). The time couse analysis of the enzyme cascade reaction of SgvM for 4-methyl-2-oxovaleric acid as a substrate starting from D- and L-methionine is shown in Figure S8.

Table S3: Retention times of substrates and methylated products in SgvM assays.

| | 4-methyl-2-oxovaleric acid | 2-oxovaleric acid |
|---|---|---|
| substrate | 18.69 min | 16.69 min |
| Methylated product | 22.29 min | 20.13 min |

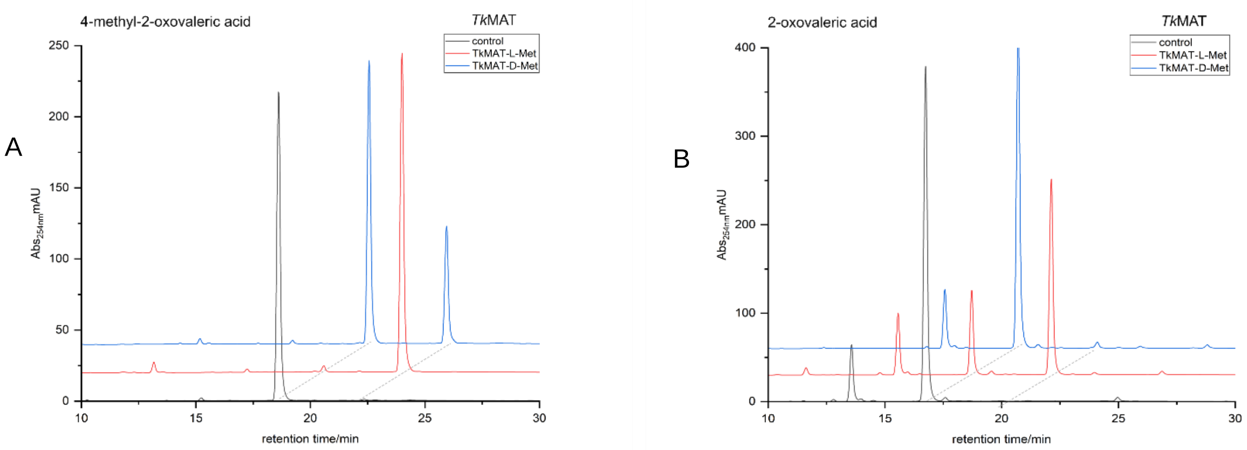

Figure S7: HPLC chromatograms of SgvM assays with different substrates. A: Reaction with 4-methyl-2-oxovaleric acid as substrate in the presence of *Tk*MAT and either L-Met, D-Met, or no enzyme (negative control). B: Reaction with 2-oxovaleric acid as substrate in the presence of *Tk*MAT and either L-Met, D-Met, or no enzyme (negative control).

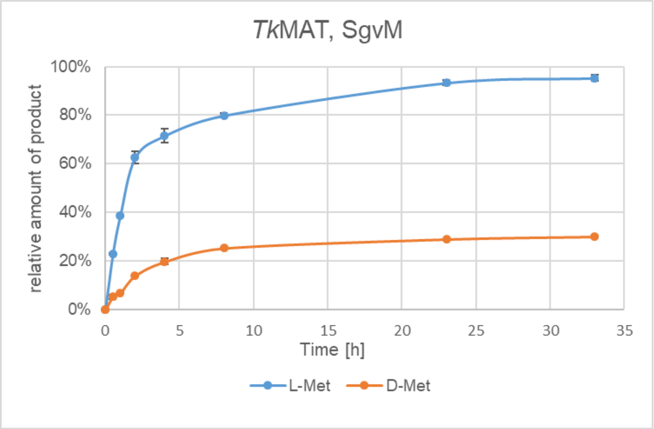

Figure S8: Time course of SgvM in three-enzyme cascade with *Tk*MAT. Error bars indicate standard deviation between triplicates. Blue: *Tk*MAT, SgvM, L-Met. Orange: *Tk*MAT, SgvM, D-Met.

##### 1.4.3 *Assay Conditions for Radical SAM Enzymes (HydG, TsrM, GenD1, QCMT) and HPLC-MS analysis*

For selected radical SAM enzyme assays, purified L-SAM and D-SAM were used instead of in situ SAM supply. To this end, SAM diastereomers were generated enzymatically with *Tk*MAT and purified by SCX chromatography followed by reversed-phase chromatography under the conditions given below (Table S4).

Table S4: Cation exchange chromatography and RP chromatography for SAM purification.

| Step | Component | Specification |
|---|---|---|
| **SCX Chromatography** | FPLC system | NGC Chromatography System (Bio-Rad Laboratories GmbH, Feldkirchen, Germany) |
| | Column | HiTrap SP FF 1 mL column (Cytiva Europe GmbH, Freiburg, Germany) |
| | Mobile phase A | 50 mM $NH_4COOH$, pH = 4.0 |
| | Mobile phase B | 700 mM $NH_4COOH$, pH = 4.0 |
| | Flow rate | 1 mL × $min^{-1}$ |
| | Gradient | 0–17 mL 0% B, 17–22 mL 0% B to 12% B, 22–29 mL 12% B to 100% B, 29–34 mL 100% B, 34–49 mL 0% B |
| | Injection volume | 10 mL |
| | Column temperature | 8 °C |
| | Detection wavelength | 260 nm |
| **RP chromatography** | HPLC system | Agilent 1260 Infinity II |
| | Column | ISERA IsaSpher 100-5 C18 AQ column, 250 mm × 8 mm (ISERA GmbH, Düren, Germany) |
| | Mobile phase A | 0.1% FA |
| | Mobile phase B | MeCN |
| | Flow rate | 1 mL × $min^{-1}$ |
| | Gradient | 0–20 min 0% B, 20–22.6 min 0% B to 30% B, 22.6–39.5 min 30% B, 39.5–41 min 30% B to 0% B, 41-60 min 0% B |
| | Injection volume | 100 µL |
| | Column temperature | 20 °C |
| | Detection wavelength | 260 nm |

*Supporting Tables and Figures (HydG, QCMT, GenD1, TsrM)*

All assays involving the radical SAM enzymes HydG, QCMT, GenD1 and TsrM were performed under strictly anaerobic conditions in an anaerobic chamber (0 ppm $O_2$, ≥2.0% $H_2$). Plasticware was transferred into the chamber at least 24 h prior to use. Buffers, water, and salt solutions were rendered anaerobic by repeated vacuum/$N_2$ cycles and were opened exclusively inside the chamber. Solid reagents were transferred into the chamber in closed reaction tubes and dissolved in anaerobic Milli-Q water immediately before use. Reactions were incubated at 37°C and 350 rpm inside the anaerobic chamber. Samples were removed from the chamber, quenched by addition of 10% (v/v) $HClO_4$ to a final concentration of 2.5% and snap-frozen in liquid nitrogen. Reactions containing GenD1, QCMT, or TsrM were protected from light during incubation.

HydG, GenD1, and TsrM were used as purified enzymes. QCMT was used in its purified, fully reconstituted form. Commercially synthesised 10-mer peptide (HFGGSQRAGV)[6] was used as QCMT substrate.

Assays probing L-SAM and D-SAM acceptance by the radical enzymes HydG, TsrM, GenD1, and QCMT were performed under the anaerobic conditions described above. In assays with in situ L-SAM supply, L-Met (2 mM), ATP (2 mM), and EcMAT (10 µM) were used; the corresponding conditions are listed in Table S5. In assays with in situ D-SAM supply, D-Met (3 mM) and TkMAT (10 µM) were used in place of the L-SAM-generating system.

Table S5: General composition of positive control radical SAM enzyme reactions performed in this study. Buffer salts and SAM supply were identical in all positive control reactions for all radical SAM enzymes. Composition of reductants and other assay compositions were adapted from literature.[2,3,7] Tris tris(hydroxymethyl)aminomethane;; Mev: methylviologen;, DTT: dithiothreitol; OHCbl: hydroxocobalamin; GentA, gentamicin A; 10-mer peptide substrate: HFGGSQRAGV; L-Trp: L-tryptophan, L-Tyr: L-tyrosine, L-Met: L-methionine

| Enzyme | Buffers and salts | SAM Supply | Reductants and additional supplements | Substrate |
|---|---|---|---|---|
| 20 µM **HydG** | | | 8 mM sodium dithionite | 0.5 mM L-Tyr |
| 15 µM **GenD1** | 50 mM Tris<br>300 mM NaCl<br>20 mM $MgCl_2$<br>50 mM KCl | 2 mM L-Met<br>2 mM ATP<br>10 µM *Ec*MAT | 1 mM Mev<br>4 mM NADPH<br>10 mM DTT<br>1 mM OHCbl | 0.4 mM GentA |
| 10 µM **QCMT** | | | 1 mM Ti(III) citrate | 50 µM 10-mer peptide substrate |
| 10 µM **TsrM** | | | 6 mM DTT<br>50 mM OHCbl | 1 mM L-Trp |

Samples were analysed by HPLC using method A, B, or C, depending on the respective enzyme assay. Assays involving TsrM, GenD1, and QCMT were analysed using HPLC method A, assays involving purified or preparative SAM were analysed using HPLC method B, and assays involving HydG were analysed using HPLC method C. The chromatographic conditions for each method are summarised in Tables S6-S8.

Table S6: HPLC-Method A.

| Component | Specification |
|---|---|
| HPLC system | Agilent 1260 infinity II system A<br>Equipped with an Agilent 1100 column compartment |
| Column | ISAsphere 100-5 C18, 250 mm × 4.6 mm, 5 µm<br>(ISERA GmbH, Germany) |
| Mobile phase A | 40 mM Na acetate, pH = 4.2 |
| Mobile phase B | MeCN |
| Flow rate | 0.5 mL × min$^{-1}$ |
| Gradient | 0–16 min 2% B to 30% B, 16–18 min 30% B,<br>18–20 min 30% to 2% B, 20–30 min 2% B |
| Injection volume | 10 µL |
| Column temperature | 26 °C |
| Detection wavelength | 260 nm, 280 nm |

Table S7: HPLC-Method B.

| Component | Specification |
|---|---|
| HPLC system | Agilent 1100 system |
| Column | ISAspher 100-5 C18 (AQ), 250 mm × 4 mm<br>(ISERA GmbH, Germany) |
| Mobile phase A | 10 mM $NH_4$FA buffer, pH = 3.5 |
| Mobile phase B | MeCN |
| Flow rate | 1 mL × min$^{-1}$ |
| Gradient | 0–12 min 0% B, 12–16 min 0% B to 20% B, 16–16.1 min<br>20% B to 30% B, 16.1–22 min 30% B,<br>22–23 min 30% B to 0% B, 23–28 min 0% B |
| Injection volume | 10 µL |
| Column temperature | 25 °C |
| Detection wavelength | 260 nm |

Table S8: HPLC-Method C.

| Component | Specification |
|---|---|
| HPLC system | Agilent 1260 infinity II system B |
| Column | ISAspher 100-5 C18, 125 mm × 4.0 mm, 3 µm, (ISERA GmbH, Germany) |
| Mobile phase A | 0.1% (V/V) formic acid |
| Mobile phase B | MeCN |
| Flow rate | 1 mL × min$^{-1}$ |
| Gradient | 0–4 min 3% B, 4-7 min 3% to 70% B, 7–7.1 min 70% to 100% B, 7.1–10 min 100% B, 10–11 min 100% to 3% B, 11–14 min 3% B |
| Injection volume | 5 µL |
| Column temperature | 26 °C |
| Detection wavelength | 254 nm, 280 nm |

*HydG Assay*

For experiments probing D-SAM acceptance by HydG, assays were performed either with in situ SAM supply or with direct addition of purified L-SAM or D-SAM. In the in situ setup, D-methionine together with *Tk*MAT was used for D-SAM generation, whereas control reactions lacking methionine were used to assess background turnover. In addition, assays were carried out with purified L-SAM or D-SAM added directly in place of the SAM-generating system. HydG turnover was monitored by HPLC method C via formation of *p*-cresol and 5′-deoxyadenosine (DOA) (Figure S9).

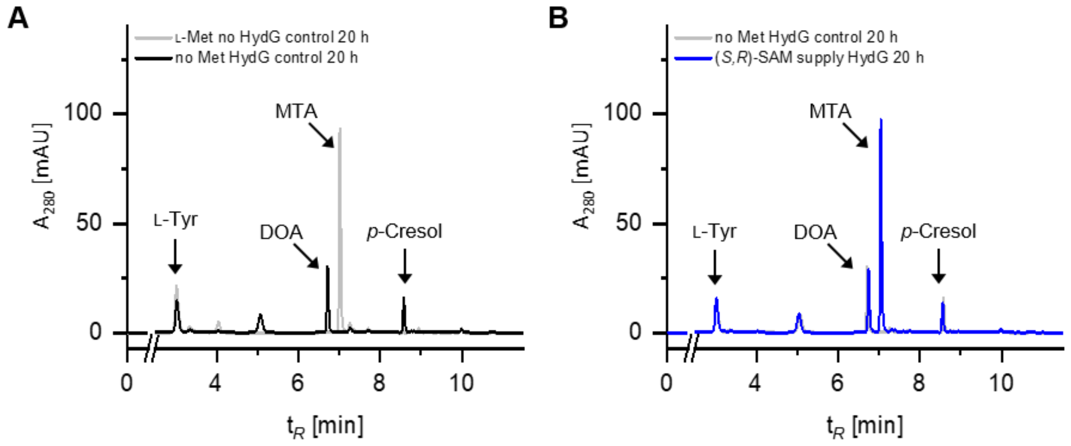

Figure S9: HPLC analysis of HydG control reactions with in situ SAM supply. (A) Comparison of a control reaction with SAM supply lacking HydG (grey) and a HydG control reaction lacking L-methionine (black). In the absence of HydG, no formation of DOA or p-cresol is observed.In contrast, when only L-methionine is omitted from an assay containing ATP, TkMAT, and HydG, formation of DOA and p-cresol is still detected. (B) Comparison of a HydG reaction with in situ SAM supply lacking methionine (grey) and a HydG reaction supplied with D-methionine (blue). Nearly identical chromatograms were obtained in both cases.

*GenD1 Assay*

For experiments probing D-SAM acceptance by GenD1, assays were performed under the general radical SAM assay conditions described above using either in situ SAM supply or direct addition of purified L-SAM or D-SAM, as indicated in the respective experiment. In assays with in situ D-SAM supply, D-Met and *Tk*MAT were used in place of the L-SAM-generating system. GenD1 turnover was monitored by HPLC method A (Table S6) via formation of SAH and 5′-deoxyadenosine (DOA) and by CE-MS-TOF via detection of DOA and GentX2. In the in situ D-SAM assays, low amounts of DOA and GentX2 were also detected in control samples lacking methionine, indicating background formation likely caused by co-purified L-methionine. In assays supplemented with purified SAM diastereomers, DOA and SAH formation was observed only with L-SAM, whereas no DOA formation was detected with D-SAM. These data indicate that GenD1 does not accept D-SAM as a cofactor.

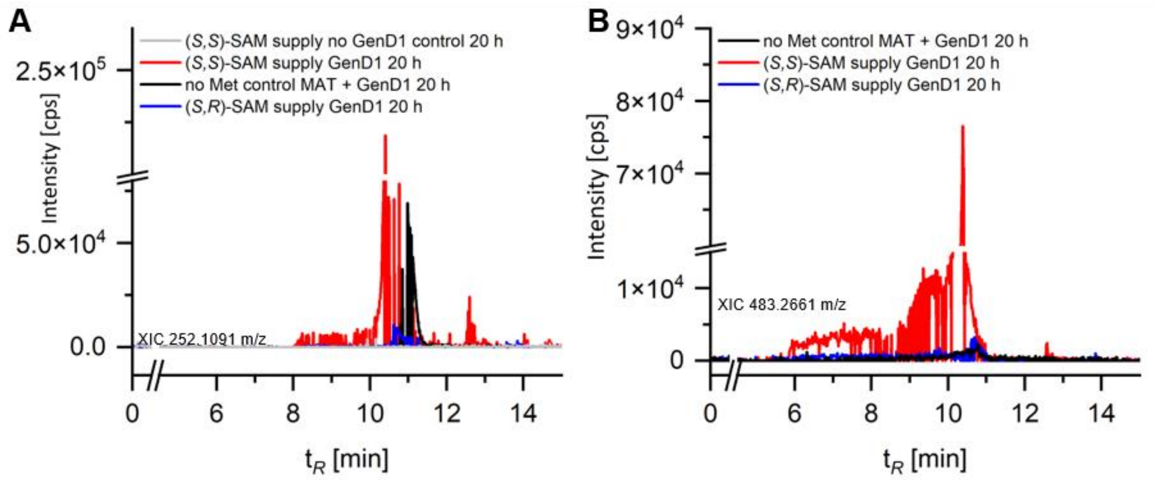

Figure S10: Extracted ion chromatograms (XICs) from CE-MS-TOF analysis of GenD1 assays. (A) XIC corresponding to the m/z value of DOA (252.1091). A GenD1 reaction with in situ D-SAM [(*S*,*R*)-SAM] supply (blue) is shown together with a control reaction without methionine (black), a control reaction with in situ L-SAM [(S,S)-SAM] supply but without GenD1 (grey), and the positive control with in situ L-SAM supply (red). Signals corresponding to DOA are observed both in the D-SAM assay and in the control lacking methionine. (B) XIC corresponding to the *m/z* value of GentX2 (483.2661). A GenD1 reaction with in situ D-SAM supply (blue) is shown together with a control reaction without methionine (black) and the positive control with in situ L-SAM supply (red). Similar signal intensities for GentX2 are observed in the D-SAM assay and in the control lacking methionine.

Samples for CE-MS-TOF analysis were centrifuged at 4 °C and 12,700 rpm for 30 min and diluted 1:1 with dd$H_2O$ prior to measurement. The mobile phase was prepared from LC-MS-grade $NH_4OH$ (25%) and formic acid. CE-MS-TOF measurements were performed on an Agilent 7100 CE instrument coupled to an Agilent 6545 Q-TOF mass spectrometer. The CE method was adapted from Moreno-González et al.[8] The corresponding instrument parameters are summarised in Table S9. Data were analysed using Agilent MassHunter software.

Table S9: MS-TOF parameters used for the analysis of aminoglycosides.

| Parameter | Specification |
| --- | --- |
| Ionisation mode | Positive |
| Gas temperature | 250 °C |
| VCap | 3500 V |
| Nebuliser gas | 10 psi |
| Drying gas | 3 L × $min^{-1}$ |
| MS-TOF Fragmentor | 100 V |
| MS-TOF Skimmer | 65 V |
| MS-TOF Oct 1 RF Vpp | 750 V |

*Supporting Tables and Figures (TsrM Assay)*

For experiments probing D-SAM acceptance by TsrM, assays were performed under the general radical SAM assay conditions described above using direct addition of purified L-SAM or D-SAM, as indicated in the respective experiment. TsrM reactions contained L-tryptophan as substrate and were analysed by HPLC method A. Formation of 2-methyltryptophan (2-MeTrp) and SAH was used to assess turnover. Product formation was observed only in reactions supplemented with L-SAM, whereas no turnover was detected with D-SAM.

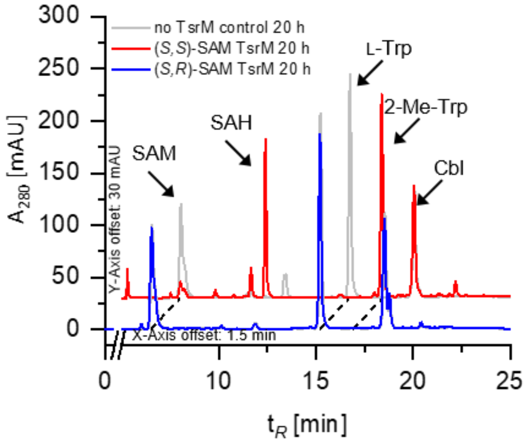

Figure S11: HPLC chromatograms of TsrM reactions supplemented with purified L-SAM [(S,S)-SAM (red) or D-SAM [(S,R)-SAM] (blue). Formation of 2-methyltryptophan (2-MeTrp) and S-adenosylhomocysteine (SAH) is observed only in the presence of L-SAM.

*QCMT Assay*

For experiments probing D-SAM acceptance by QCMT, a synthetic 10-mer peptide (HFGGSQRAGV) was used as substrate. In the D-SAM in situ supply cascade using *Tk*MAT, formation of the methylated peptide was detected by LC-MS. This result was confirmed using purified D-SAM, where peaks corresponding to the methylated peptide were observed in extracted ion chromatograms and were absent in control reactions lacking SAM or QCMT. These data show that D-SAM is accepted as a cofactor by QCMT, albeit with low turnover.

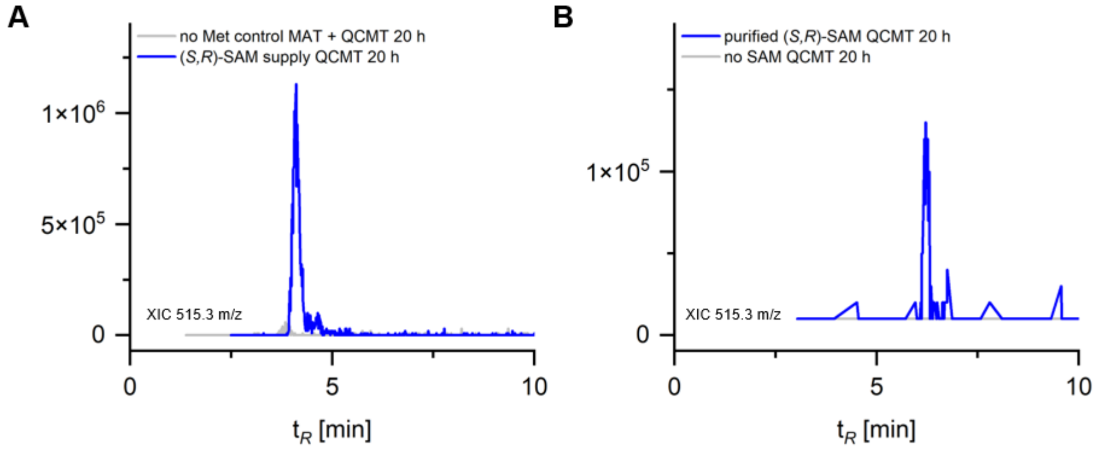

Figure S12: XICs from LC-MS analysis of QCMT reactions. The extracted ion chromatograms correspond to the expected m/z value of the methylated 10-mer peptide (515.3). (A) QCMT reaction with in situ D-SAM ((*S*,*R*,)-SAM) supply (blue) compared to a control reaction without Met (grey) (B) QCMT reaction supplemented with purified D-SAM (blue) compared to a control reaction without added SAM (grey). The methylated peptide is detected only in the sample containing D-SAM.

LC-MS method used for analysis of QCMT assay samples. Components and specifications of the chromatographic and mass spectrometric conditions are summarised in Table S10.

Table S10: Components and specifications of LC-MS method for QCMT assay.

| Component | Specification |
|---|---|
| HPLC system | Sciex Exion LC AC system |
| Column | ISAspher C4 300 Å – 5 µm 150 mm × 2.0 mm (ISERA GmbH, Düren, Germany) |
| Mobile phase A | 0.1% formic acid in $ddH_2O$ |
| Mobile phase B | 0.1% formic acid in MeCN |
| Flow rate | 0.2 mL × $min^{-1}$ |
| Gradient | 0–1 min 10% B, 1–8 min 10% B to 70% B, 8–15 min 70% B to 90% B, 15–23 min 90% B, 23–40 min 10% B |
| Injection volume | 5 µL |
| Column temperature | 25 °C |
| Mass spectrometer | Sciex 4500 |
| Ionisation mode | Positive |
| Temperature | 200 °C |
| Ion spray voltage | 4500 V |
| Declustering potential | 100 V |
| Entrance potential | 10 V |
| Curtain gas | 25 psi |
| Nebuliser gas | 40 psi |
| Heater gas | 50 psi |

*Calculation of QCMT conversion*

Conversion was calculated from the AUC values of the unmethylated peptide (Pep) and the methylated peptide (MePep) in the extracted ion chromatograms according to

$$Conversion\ [\%] = \frac{AUC(MePep)}{\mathrm{AUC}(Pep) + AUC\ (MePep)}\ x\ 100$$

For L-SAM, Pep/MePep AUC values of 5231.5/1015107, 11344/1069677.5, and 8989.5/1193567 gave conversions of 99.49%, 98.95%, and 99.25% (conversion >99%). For D-SAM, Pep/MePep AUC values of 404243/11004, 389123/16775, and 353912/12880.5 gave conversions of 2.65%, 4.13%, and 3.51% (average 3.43%). No methylated peptide was detected in the no SAM controls.

##### 1.4.4 *Assay Conditions for Cyclase PaAzeJ Synthase and HPLC-MS Analysis*

The three-enzyme cascade was conducted with 3.0 mM L- or D-methionine, 3.0 mM ATP, 3.5 µM *Pa*AzeJ, 10 µM *Uu*MAT, and 1.0 µM *Ec*MTAN in 50 mM MOPS buffer (pH 7.5) with 50 mM KCl and 20 mM $MgCl_2$ in a 600 µL reaction scale. Assays were performed at 30 °C and 350 rpm in an Eppendorf Thermomixer. Samples were taken at specified time points and directly transferred to a centrifugal filter device (Vivaspin 500, 10 kDa MW cutoff, Sartorius) to remove enzymes from the reaction. For the derivatisation 50 µL sample were added to 20 µL 1 M $NH_4HCO_3$ followed by 20 µL 1-Fluoro-2,4-dinitrophenyl-5-L-alaninamide (1% in acetone). Samples were then incubated at 40 °C, 700 rpm for 1h before adding 20 µL HCl (2 M). After centrifuging at 4°C, 12,700 rpm (Eppendorf centrifuge 5427 R; Eppendorf, Hamburg, Germany) for 1 h, samples were transferred to HPLC glass vials (Isera, Germany) and subjected to HPLC analysis.

HPLC analysis was carried out on an Agilent 1100 system equipped with an ISAspher 100–5 C18 column (250 mm, 4 mm, 5 µm). A gradient method was used with 1 mL/min flowrate and 1% acetonitrile (MeCN) as well as 0.1% formic acid (v/v) in water as eluent A and MeCN as eluent B. The gradient was as follows: 0-7 min 90% → 40% A, 7-12 min 40% → 5% A, 12-15 min 5% → 90% A, 15-17.5 min 90% A. The detection wavelength was set to 348 nm and the injection volume was 10 µL.

Derivatisation with 4-fold molar excess of Marfey's reagent yielded the expected diastereomers, with the (L, L) derivative formed from L-SAM and the (D,L) derivative from D-SAM. These diastereomers were clearly distinguished by HPLC. This method was described by Yan et al (2019).[9]

Retention times of derivatised products and standards are summarised in Table S11, enabling assignment of D- and L-azetidine-2-carboxylic acid (D-AZC and L-AZC, respectively).

Table S11: Retention times of substances from *Pa*AzeJ assay.

| ***Substance*** | ***Retention time [min]*** |
|---|---|
| *D-AZC (derivatised)* | *8.05* |
| *L-AZC (derivatised)* | *8.31* |
| *Marfey's reagent* | *9.30* |
| *L-Methionine (derivatised)* | *9.42* |
| *D-Methionine (derivatised)* | *10.00* |

The products were analysed by HPLC, and the resulting chromatograms are shown in Figure S13 for the (L,L)-Marfey derivative and Figure S14 for the (D,L)-Marfey derivative.

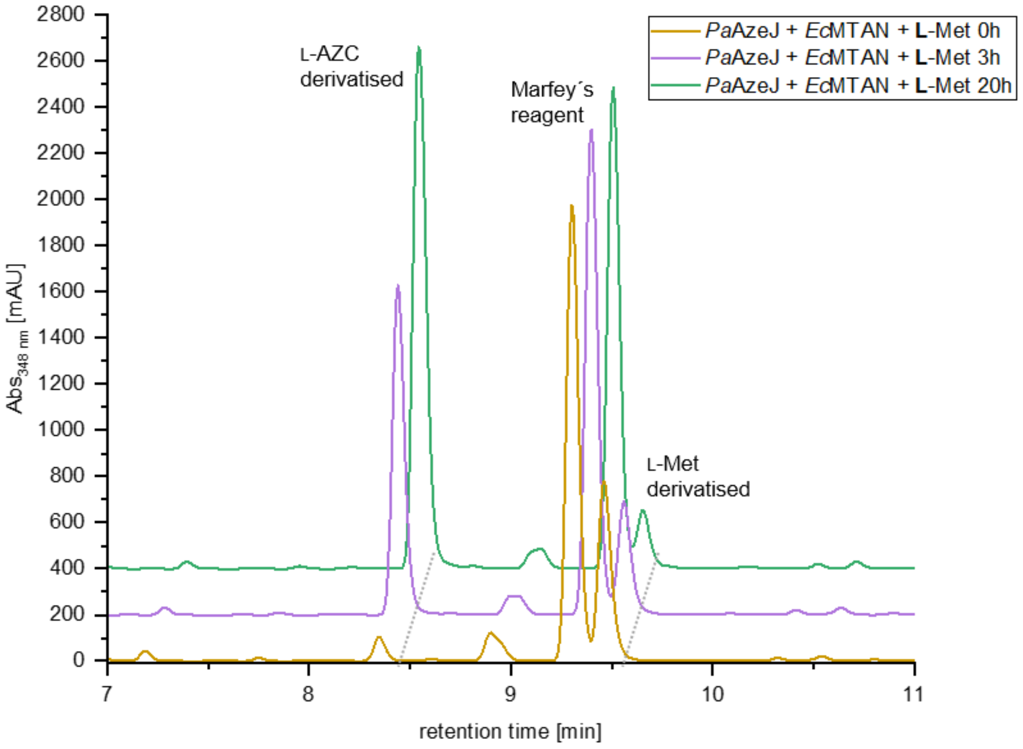

Figure S13: HPLC chromatograms of derivatised products from AzeJ-catalysed reactions. (L,L)-Marfey derivatives obtained from L-methionine (via L-SAM).

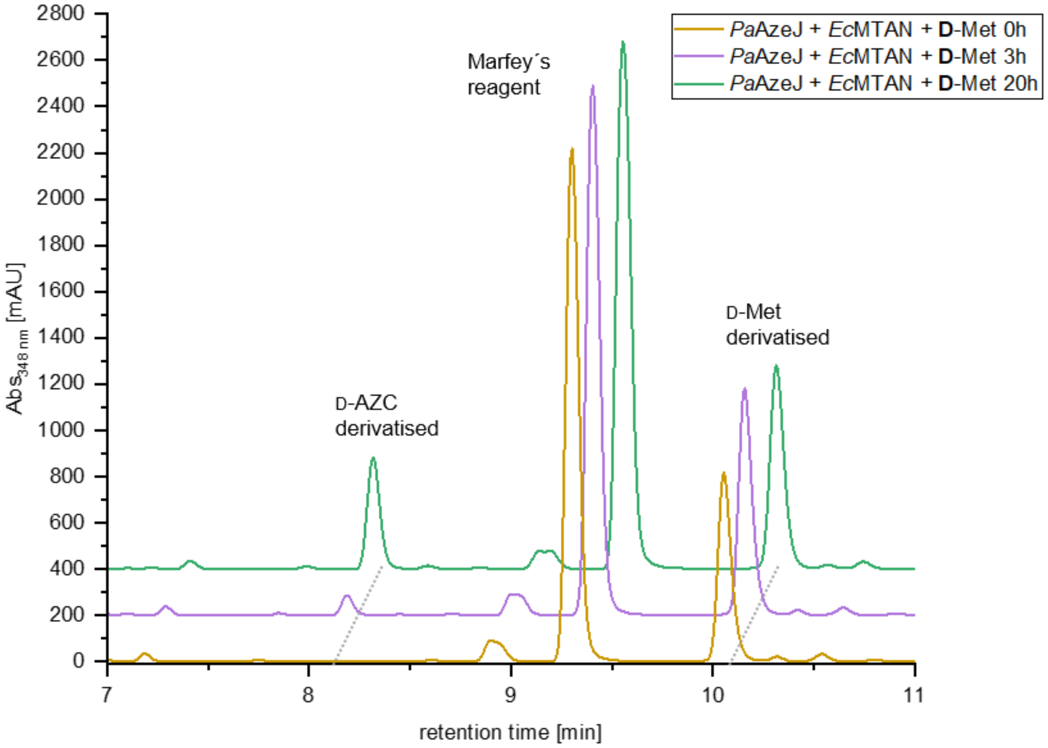

Figure S14: HPLC chromatograms of derivatised products from *Pa*AzeJ-catalysed reactions. (D,L)-Marfey derivatives obtained from D-Met (via D-SAM).

##### 1.4.5 Assay Conditions for Cyclase ACC Synthase and HPLC-MS/MS Analysis

The enzyme ACCS (10 µm) and either L-SAM or D-SAM (1 mm) were incubated in reaction buffer (50 mM HEPES, 150 mM NaCl, pH 8.5) at 30 °C for 5 h in a total reaction volume of 100 µL. Reactions were quenched by the addition of 100 µL methanol. Negative controls were performed in the absence of enzyme. For improved mass spectrometric detection, 50 µL of the assay mixture was combined with 50 µL of sat. $NaHCO_3$ solution and 12.5 µL of dansyl chloride (20 mM in ethanol). Following, the reaction was incubated for 1 h at 37 °C. Samples were subsequently analysed by HPLC-MS/MS.

HPLC-MS/MS measurement was performed on a Bruker Impact II coupled with a Bruker Elute UHPLC system. The reaction product was detected in positive ionisation mode with MS parameters: nitrogen as dry gas at 7.5 L/min at 2.5 bar, capillary temperature of 220 °C, capillary voltage of 4500 V and a collision energy of 8 eV. Liquid chromatography was conducted using an EC 100/2 NUCLEODUR HILIC column (Machery Nagel, 5 µm particle size) with a temperature set point of 40 °C. The mobile phases consisted of solvent A: 5 mM ammonium formate (pH 7.0), and solvent B: MeCN. The gradient was: 0.00–2.00 min, 98% B (isocratic, 0.25 mL/min); 2.00–8.00 min, 98% to 90% B (0.25 mL/min); 8.00–9.00 min, 90% to 40% B (0.25 mL/min); 9.00–23.00 min, 40% B (isocratic, 0.50 mL/min); 23.00–23.50 min, 40% to 98% B (0.50 mL/min); 23.50–24.50 min, 98% B (isocratic, 0.25 mL/min). Injection volumes were 5 µL.

*Supporting Tables and Figures (ACCS)*

Figure S15a shows the HPLC-MS chromatograms of the ACCS assays. Figure S15b displays the MS/MS spectra used for product verification.

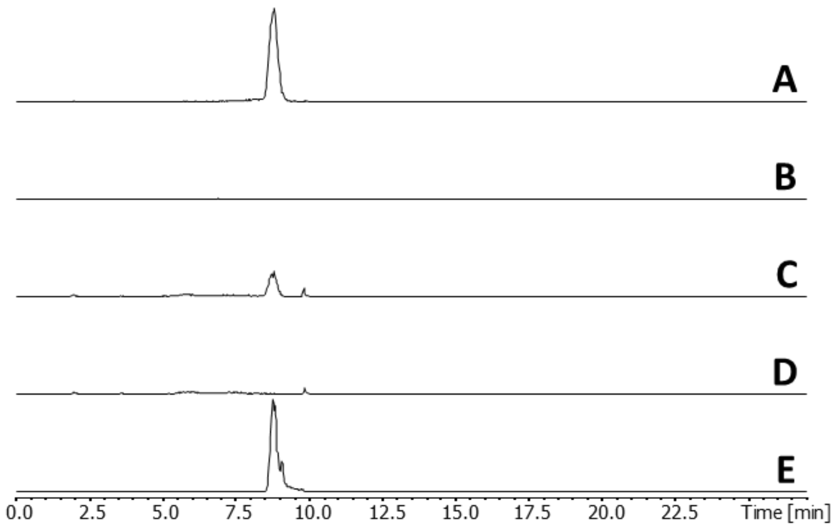

Figure S15a: HPLC-MS chromatograms after derivatisation with dansyl chloride. Extracted ion chromatograms for dansylated product ACC is shown (335.1057 *m*/*z*) A: L-SAM with ACCS; B: corresponding negative control of A; C: D-SAM with ACCS; D: corresponding negative control of C; E: dansylated ACC reference.

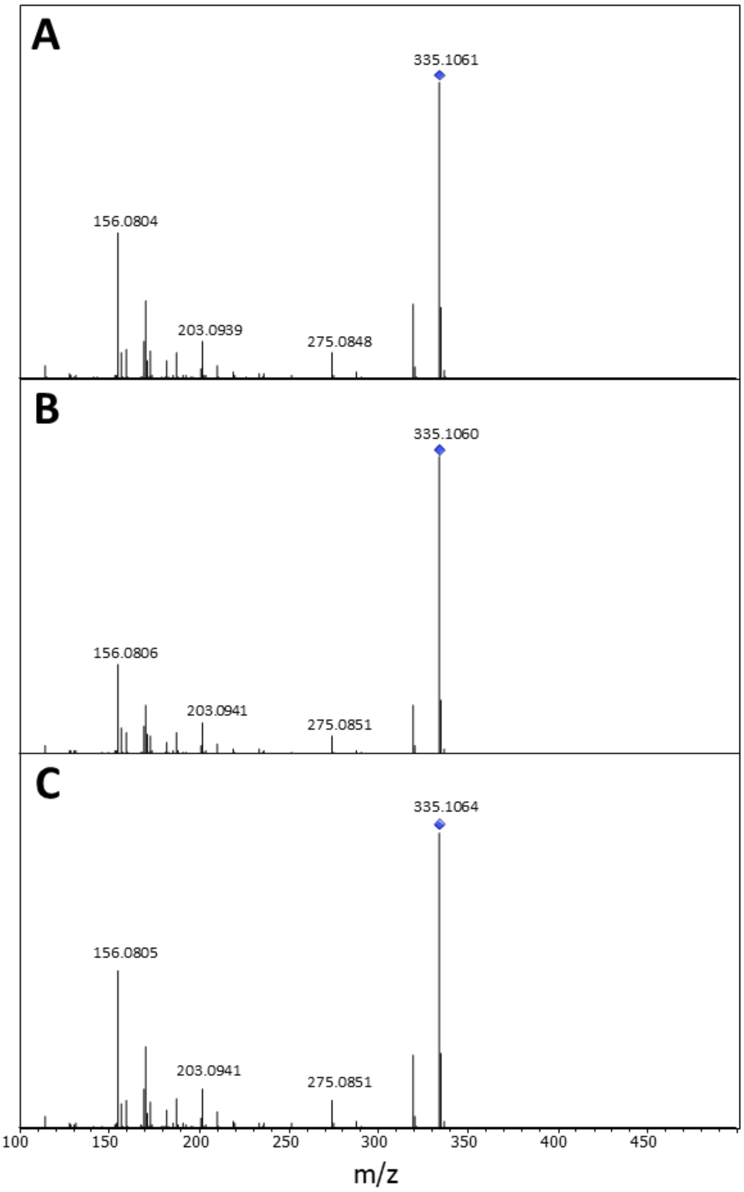

Figure S15b: MS/MS spectra acquired in positive ion mode. The enzymatic reactions and the reference were derivatised with dansyl chloride. A: L-SAM with ACCS; B: D-SAM with ACCS; C: ACC reference

#### 1.5 Docking and Structural Analysis

Molecular docking simulations were carried out based on crystal structures of enzymes in complex with SAH or SAM analogues retrieved from the Protein Data Bank (PDB)[10]. Docking of L-SAM and the according enzyme substrates was predominantly performed using the co-folding approach of the web-based Chai-1[11] algorithm. For the predictions, amino acid sequences given in the respective PDB entries were used together with the SMILES codes of the small molecules retrieved from PubChem. The option 'Use MSAs' (MMseqs2) as well as 'Use Templates' were activated. Docking of D-SAM and D-configured small molecules was performed using the web-based CB-Dock2[12] algorithm. Here, the according '.pdb-files' retrieved from the PDB were uploaded together with the '.sdf-files' of the three-dimensional small molecule structures retrieved from PubChem. Docking was then performed using the cavity search function. Finally, the resulting models were visualised and analysed with the PyMOL[13] Molecular Graphics System, Version 3.1.0 to evaluate binding conformations and key interactions. More detailed information regarding the performed molecular docking experiments, including the obtained scores, are given in Table S12.

Table S12: Molecular docking experiments.

| Docking experiment | PDB template | Used software | Docking score |
|---|---|---|---|
| 4,5,7-trihydroxy-3-phenylcoumarin and L-SAM in CouO | 5M58 (Sequence) | Chai-1 | 0.86 (ptm)/0.83 (iptm) |
| D-SAM in CouO | 5M58 | CB Dock2 | -8.3 |
| D-SAM in SgvM | 8FTS | CB Dock2 | -9.3 |
| L-SAM and ketoleucine in SgvM | 8FTS (Sequence) | Chai-1 | 0.95 (pTM) / 0.92 (iptm) |
| D-SAM in AzeJ | 8RYE | CB Dock2 | -9.5 |
| D-SAM in AzeJ (wrong) | 8RYE | CB Dock2 | -9.6 |
| D-AZC in AzeJ | 8RYE | CB Dock2 | -4.4 |
| L-SAM in AzeJ | 8RYE (Sequence) | Chai-1 | 0.92 (pTM) / 0.87 (iptm) |
| L-SAM in ACCS | 1M4N | CB Dock2 | -2.7 |
| D-SAM in ACCS | 1M4N | CB Dock2 | -2.6 |

##### *1.5.1 C-Methyltransferases: Structural Modelling and Docking of CouO and SgvM*

Figures S16 and S17 present the docked conformations of both SAM diastereomers within the active sites of CouO and SgvM, respectively, based on available crystal structures (PDB: 5M58 for CouO[14] and PDB: 8FTS for SgvM[15]). Figures S18 and S19 provide a comparative analysis of key geometric parameters relevant to methyl group orientation and potential $S_N2$ transition state alignment in the presence of either L- or D-SAM.

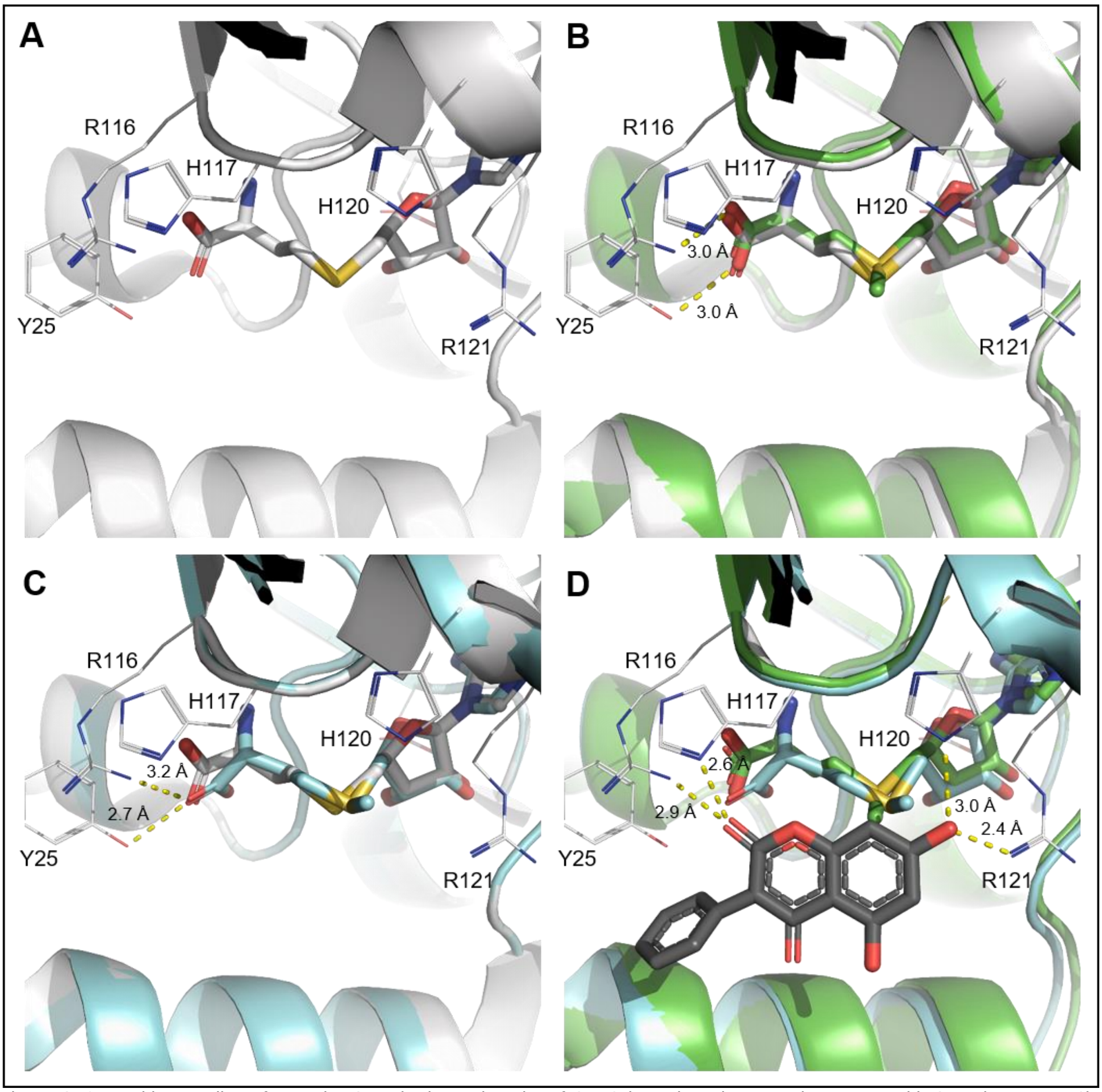

Figure S16: Docking studies of L- and D-SAM in the active site of CouO based on the crystal structure with SAH (PDB: 5M58). While the SAM diastereomers are shown as sticks and important active site residues are shown as white lines, likely interactions between the active side residues and ligands are depicted as yellow dashed lines. A: Crystal structure of CouO with bound SAH used as the template structure. B: Docking pose of L-SAM into the active site of CouO. C: Docking pose of D-SAM into the active site of CouO. D: Superposition of docked L-SAM and D-SAM together with the substrate 4,5,7-trihydroxy-3-phenylcoumarin (dark grey sticks) in the active site of CouO.

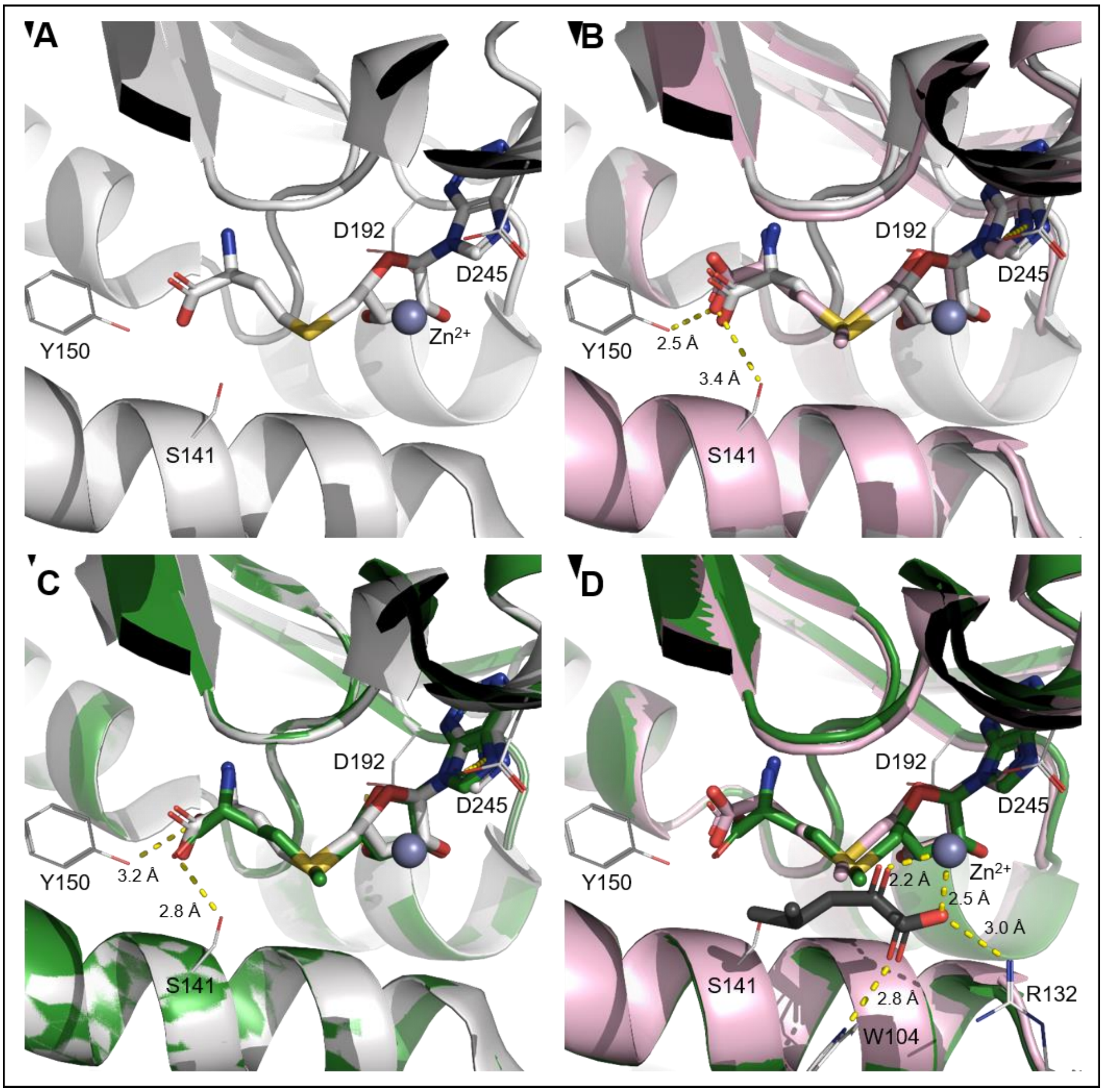

Figure S17: Docking studies of L- and D-SAM in the active site of SgvM based on the crystal structure with SAH, α-ketoleucine and $Zn^{2+}$ (PDB: 8FTS). While the SAM diastereomers are shown as sticks and important active site residues are shown as white lines, likely interactions between the active side residues and ligands are depicted as yellow dashed lines. A: Crystal structure of SgvM with bound SAH and $Zn^{2+}$ (grey sphere) used as the template structure. B: Docking pose of L-SAM in the active site of SgvM. C: Docking pose of D-SAM in the active site of SgvM. D: Superposition of docked L-SAM and D-SAM together with the substrate 4-methyl-2-oxovalerate (dark grey sticks) within the active site.

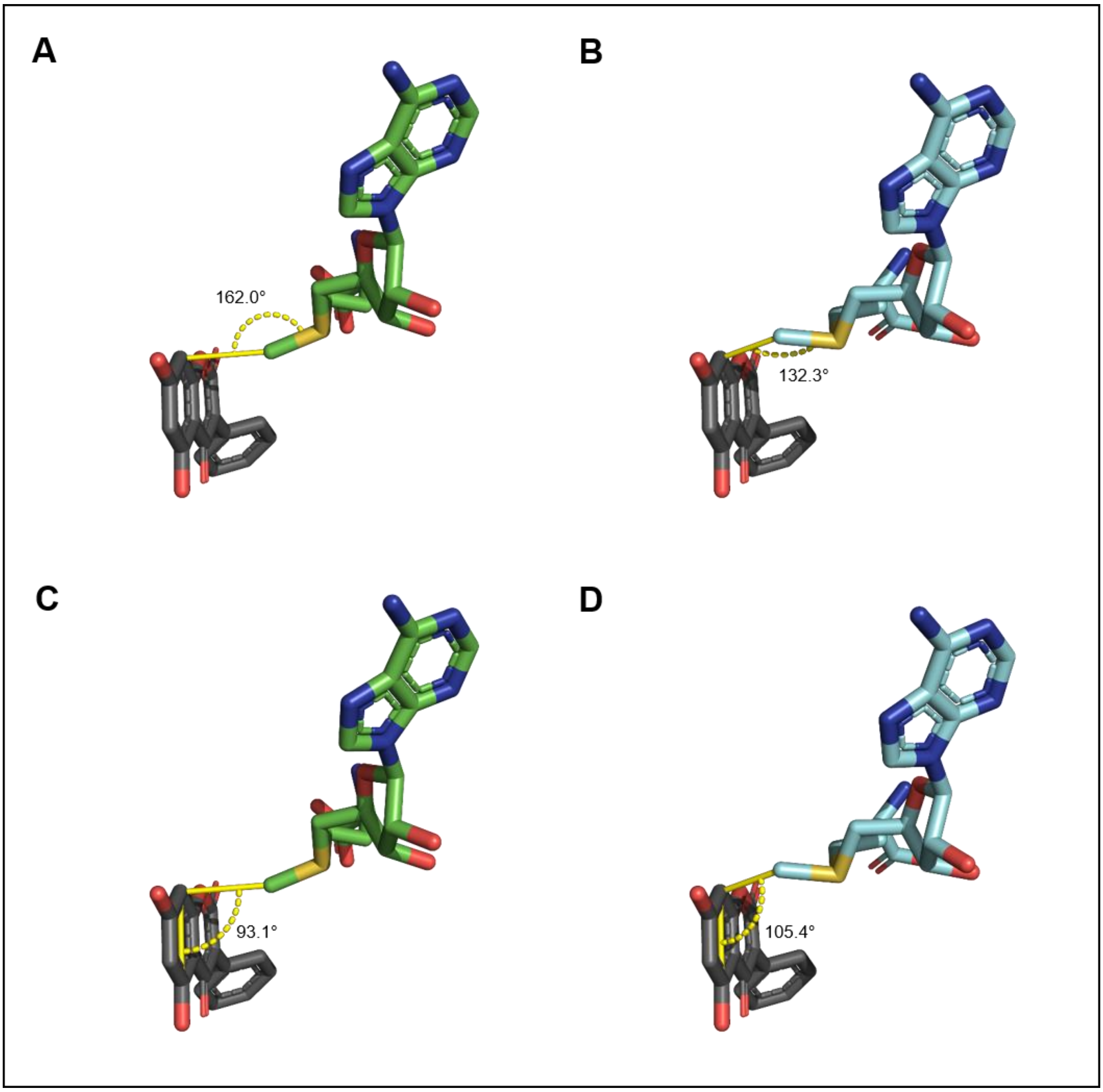

Figure S18: Comparison of active site geometries in CouO with L- and D-SAM in complex with the native substrate 4,5,7-trihydroxy-3-phenylcoumarin. The small molecules are shown as sticks while angle measurements are indicated by yellow (dashed) lines. A: Binding pose with L-SAM, showing a near-linear $S_N2$-like geometry with an angle of 162.0° between the nucleophilic carbon of the substrate, the methyl group, and the sulfonium sulfur. B: Binding pose with D-SAM, displaying a reduced angle of 132.3°, indicating a deviation from ideal methyl transfer geometry. C: Torsional alignment of L-SAM in the active site with an angle of 93.1  between the methyl group and the substrate's aromatic plane. D: Corresponding pose with D-SAM showing a larger angle (105.4°), suggesting suboptimal orientation for methyl transfer.

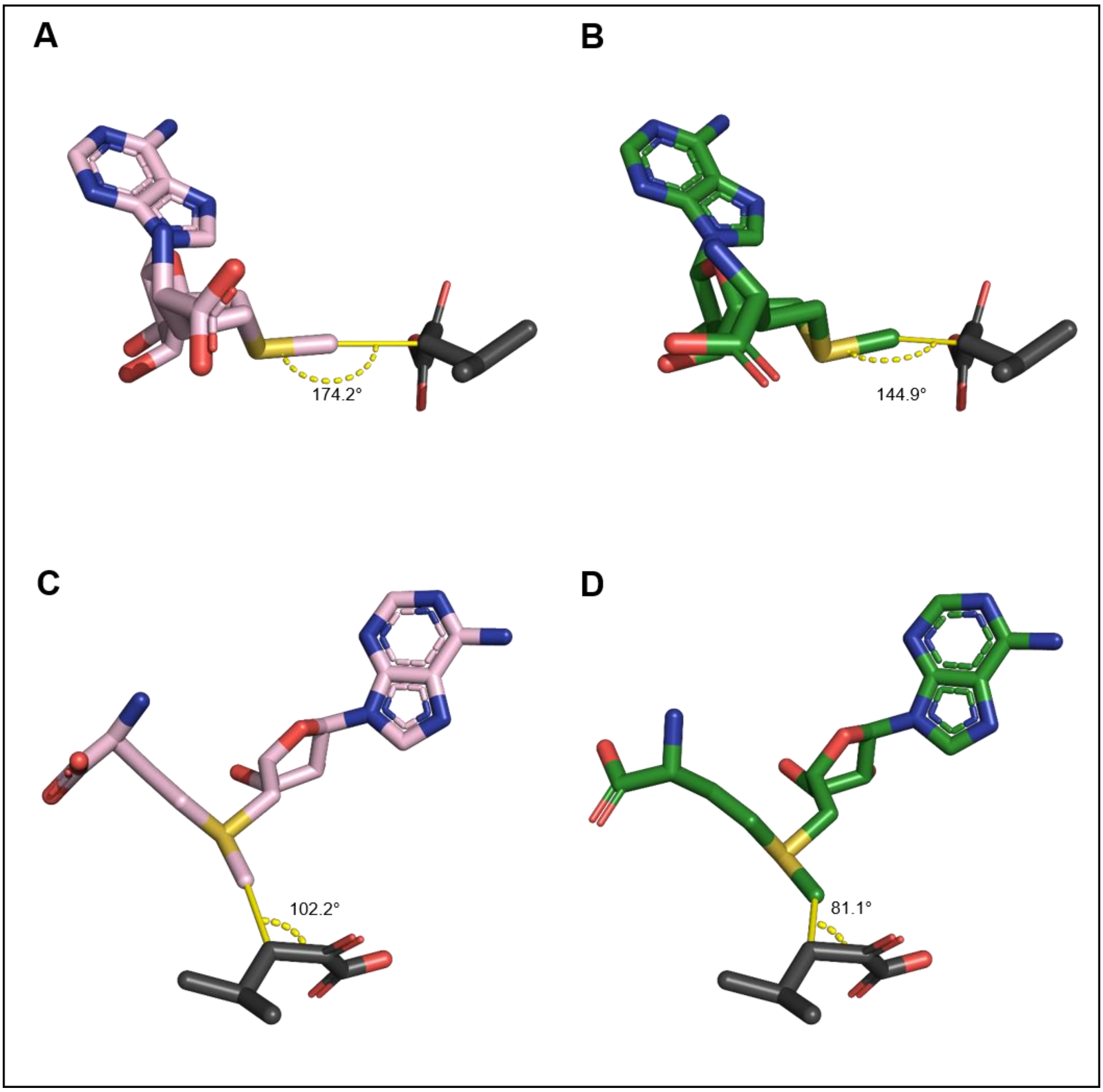

Figure S19: Comparison of active site geometries in SgvM with L- and D-SAM in complex with 4-methyl-2-oxovalerate. The small molecules are shown as sticks while angle measurements are indicated by yellow (dashed) lines. A**:** Binding pose with L-SAM, showing a favourable $S_N2$-like geometry with an angle of 174.2° between the nucleophilic carbon, the methyl group, and the sulfonium sulfur. B**:** Binding pose with D-SAM, displaying a reduced angle of 144.9°, indicative of a distorted geometry unfavourable for methyl transfer. C: Orientation of the methyl group in L-SAM relative to the molecular plane of the substrate, with an observed angle of 102.2°. D**:** Orientation of the methyl group in D-SAM relative to the molecular plane of the substrate, with an observed angle of 81.1°.

##### *1.5.2 Cyclases: Structural Modelling and Docking of AzeJ*

Figure S20 presents the docked conformations of both SAM diastereomers within the active site of AzeJ, based on an available crystal structures (PDB: 8RYE in complex with MTA and L-AZC)[16]. Particular attention was given to the distance between the Cγ atom and the amino group, as this position is involved in the cyclisation reaction.

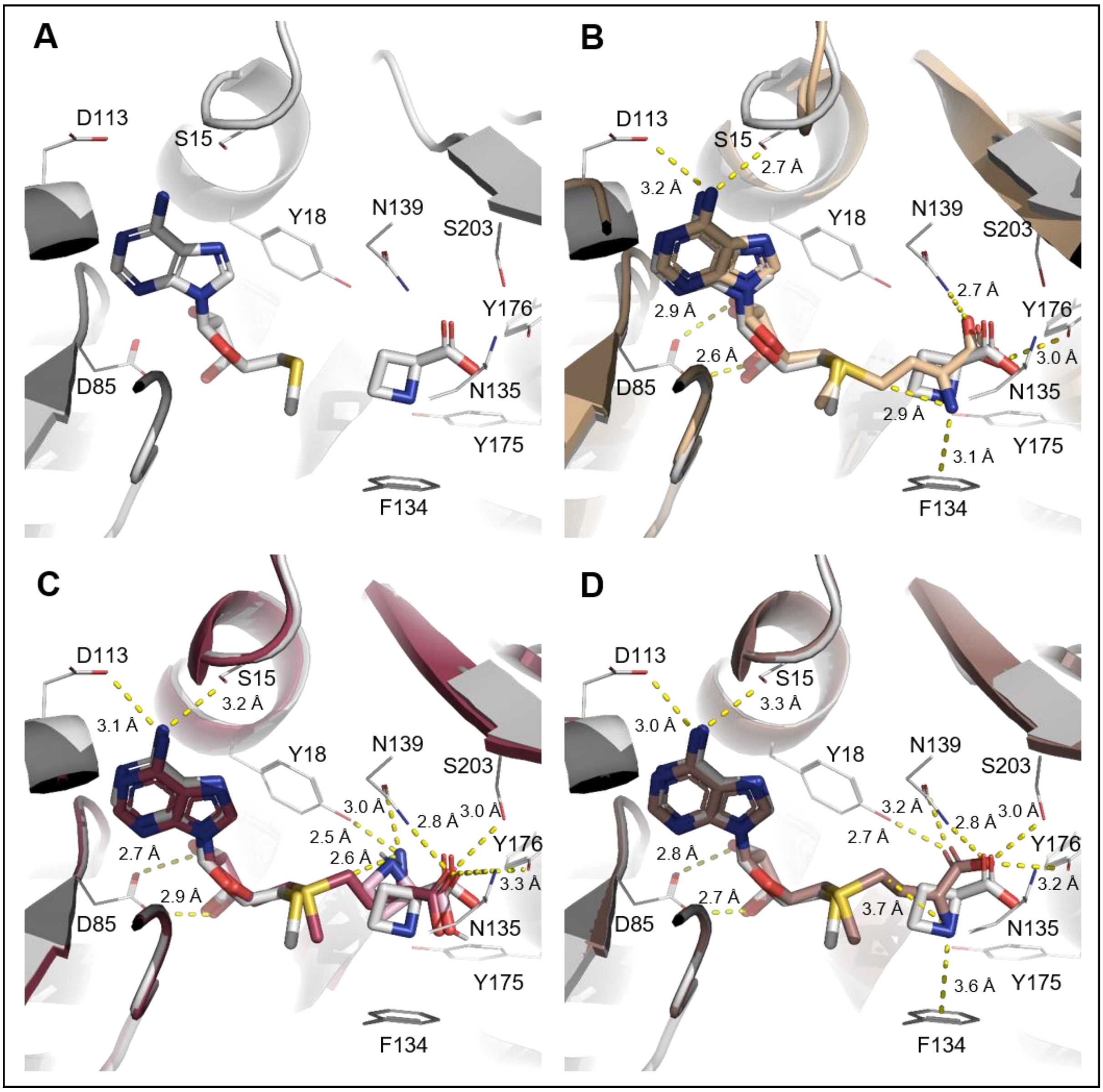

Figure S20: Structural comparison of L- and D-SAM binding modes in AzeJ based on crystal structure and docking analysis. While the SAM diastereomers are shown as sticks and important active site residues are shown as white lines, likely interactions between the active side residues and ligands are depicted as yellow dashed lines. A: Crystal structure of AzeJ in complex with L-AZC and MTA (PDB: 8RYE). B: Docked conformation of L-SAM (with MTA) and L-AZC, showing a favourable distance of 2.9 Å between the Cγ atom and the amino group. C: Docked conformations of D-SAM (with MTA) and D-AZC, showing a distance of 2.6 Å between the Cγ atom and the amino group; however, no stabilising interaction with Phe134 is observed. D: Alternative binding pose of D-SAM in which the amino group could be positioned for activation by Phe134; however, the distance and the angle between the Cγ atom and the amino group are unfavourable.

*1.5.3 Cyclases: Structural Modelling and Docking of ACCS*

Figure S21 illustrates the docking experiments performed with L- and D-SAM in the active site of ACCS. The crystal structure of ACCS in complex with an amino oxy analogue (AMA) and the cofactor PLP (PDB: 1M4N) was used as template for SAM docking. The crystal structure of ACCS in complex with PLP (PDB: 1IAX; dark grey) captures the enzyme in its dimeric state. It was superimposed to visualise the interactions of SAM with the adjacent monomer as well as the positioning of PLP without a covalently bound substrate molecule. Particular attention was given to the distance between the amino group of K273 and the α-carbon atom of SAM at which the C–H deprotonation takes place.

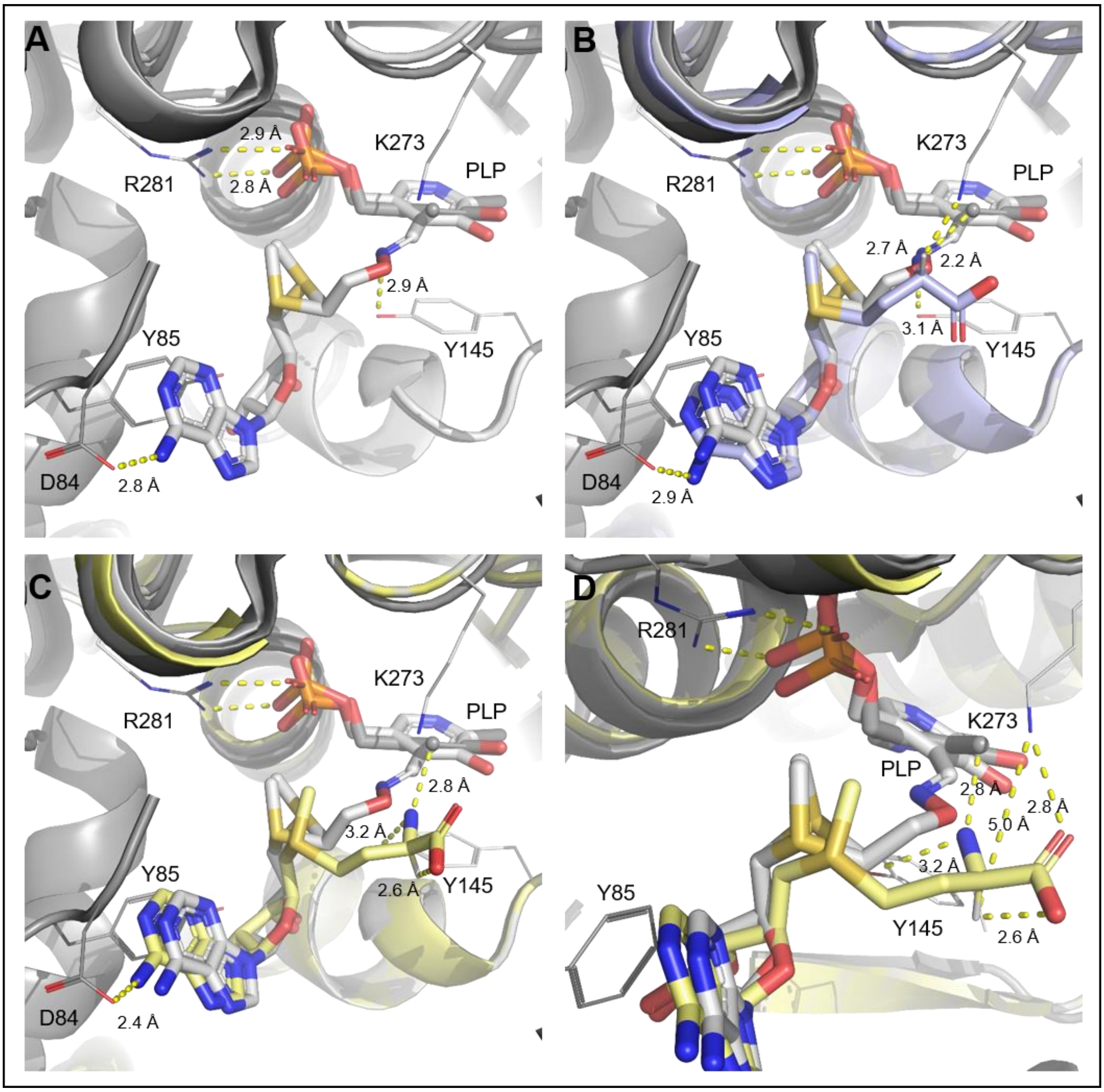

Figure S21: Dimeric structure and SAM binding modes of ACCS. Both the substrate analogue AMA and cofactor PLP are shown as sticks, while the protein backbone is depicted in cartoon representation, with key residues highlighted in white lines. The sulfonium site of AMA is shown in two different configurations, caused by racemisation during crystallisation, according to the original publication.[17] Likely interactions and distances are indicated with yellow dashed lines. A: Crystal structure of ACCS (PDB: 1M4N) in complex with AMA and PLP, shown alongside the PLP bound ACCS dimer (PDB: 1IAX, dark grey) for comparison. B. Docked conformation of L-SAM showing a favourable distance of 2.7 Å between the C-H atom at the α-carbon of L-SAM and the amino group of K273. C: Docked conformation of D-SAM, in which the C-H group of D-SAM points away from the amino group of K273, preventing direct deprotonation. D: Close-up view of D-SAM, highlighting a 2.6 Å distance between C-H atom and the carboxylate group (sufficient for intramolecular deprotonation) and a 2.8 Å distance from the carboxyl group to K273 (sufficient for deprotonation).

##### 1.6 Summary of Reported and Observed D-SAM Acceptance by SAM-dependent enzymes

To compare the results obtained in this study with previous reports, Table S13 summarises examples of SAM-dependent enzymes tested with D-SAM as cofactor or substrate. Where available, the reported extent of conversion is indicated.

Table S13: Examples of SAM-dependent enzymes reported to accept D-SAM as cofactor or substrate.

| **Enzyme (organism)** | **D-SAM as cofactor/ substrate (conversion)** | **Tested acceptor substrate** | **Reference** |
|---|---|---|---|
| anthranilate *N*-methyl-transferase, ANMT (*Ruta graveolens*) | Yes (>99%) | 2-amino-4-nitrophenol | this work |
| caffeate *O*-methyl-transferase, CaOMT (*Prunus persica*) | Yes (92%) | 2-amino-4-nitrophenol | this work |
| *C*-methyltransferase SgvM (*Streptomyces griseoviridis)* | Yes (29%) | 4-methyl-2-oxovalerate | this work |
| *C*-methyltransferase CouO (*Streptomyces rishiriensis)* | No | 4,5,7-trihydroxy-3-phenylcoumarin | this work |
| glutamine C-methyl-transferase QCMT (*Methanoculleus thermophiles)* | Yes (3%) | 10-mer peptide (HFGGSQRAGV) | this work |
| cobalamin-dependent radical SAM methyltransferase GenD1 (*Micromonospora echinospora)* | No | Gentamicin A | this work |
| tyrosine lyase HydG (*Thermoanaerobacter italicus)* | No | L-tyrosine | this work |

| | | | |
|---|---|---|---|
| tryptophan 2-*C*-methyltransferaseTsrM (*Streptomyces laurentii)* | No | L-tryptophan | this work |
| azetidine-2-carboxylic acid synthase, AzeJ (*Pseudomonas aeruginosa or Saccharotrix sp. NRRL B-16348)* | Yes (>30% for *Pa*AzeJ or >20% for *Sac*AzeJ) | SAM | this work |
| engineered 4-O-methyl-transferase (*Eriobotyra japonica)* | Yes (20%) | ferulic acid | Turner et al.[18] |
| 1-aminocyclopropyl-1-carboxylic acid synthase (ACCS) (*Malus domestica)* | Yes (traces) | SAM | this work |
| *N*-acetylserotonin *O*-methyl-transferase (*bovine pineal glands)* | No | *N*-acetyl-serotonine | Nakamura and Schlenk[19] |
| L-homocysteine *S*-methyl-transferase from (*Saccharomyces cerevisiae)* | Yes (>20%) | L-homocysteine | Nakamura and Schlenk[19] |
| histamine *N*-methyl-transferase (*pig brain)* | Yes | histamine | Nakamura and Schlenk[19] |
| guanidinoacetate *N*-methyl-transferase (*pig liver)* | Yes (>5% conversion) | guanidineacetate | Nakamura and Schlenk[19] |

#### 1.7 Production, crystallisation, and structure determination of *Sac*Azej

##### *1.7.1 Production of recombinant SacAzeJ*

The gene encoding AzeJ from *Saccharotrix sp. NRRL B-16348* was cloned into the first multiple cloning site of a modified pCOLA-DUET1 vector, encoding additional *N*-terminal MBP, Strep-tag II and a TEV protease cleavage site. The protein was expressed in E. coli BL21 (DE3) Star in ZYM-5052 auto-inducing medium[20] at 20 °C for 20–24 h. The cell pellet was resuspended in a buffer containing 20 mM Tris/HCl pH 8.5, 300 mM NaCl, 10% (v/v) glycerol, one tablet of Complete EDTA-free protease inhibitor cocktail (Roche), and lysed by sonication. The protein was isolated from the supernatant after centrifugation for 1 h at 100,000 × g using a self-packed 10 mL column with Strep-Tactin Superflow High-Capacity resin (IBA) and eluted from the column with a single step of 5 mM D-desthiobiotin. Affinity and solubility tags were removed by cleavage with TEV protease (1:30) over night at 4 °C. Gel filtration was carried out using a HiLoad 16/600 Superdex 75 pg column (GE Healthcare/Cytiva) in 20 mM Tris/HCl pH 8.5, 300 mM NaCl, 10% (v/v) glycerol. All chromatography steps were carried out using an Äkta Purifier system (GE Healthcare). The peak fractions of the size exclusion were analysed by SDS-PAGE, concentrated, flash-frozen in liquid nitrogen, and stored at -80 °C until crystallisation screening.

##### *1.7.2 Crystallisation*

Crystallisation trials were set up at room temperature with a Crystal Gryphon crystallisation robot (Art Robbins Instruments) in Intelli 96-3 plates (Art Robbins Instruments) with 200 nL protein solution and 200 nL reservoir solution. Crystals were obtained at room temperature in condition E4 of the JCSG Core II screen (QIAGEN) at 6 mg/mL protein. Optimised crystals of sufficient quality for data collection were obtained in 1.1 M LiCl and 0.1 M Tris/HCl pH 7.0. Crystals were cryoprotected with 10% (v/v) (*R*,*R*)-2,3-butanediol and, after harvesting, flash-cooled and stored in liquid nitrogen until measurement.

##### *1.7.3 Data collection and processing*

Data collection was performed at beamline P11 at the PETRA III storage ring (Deutsches Elektronen-Synchrotron, Hamburg, Germany)[21]. All datasets were recorded at a temperature of 100 K. Data processing was carried out using the autoPROC[22] toolbox (Global Phasing), executing XDS[23], Pointless[24], and Aimless.[25]

##### *1.7.4 Structure determination, refinement, and model building*

The structure of SacAzeJ was determined by molecular replacement using a model generated by AlphaFold2 as a search model for Phaser[26] from the Phenix suite[27]. The structural model was built using Coot[28], and crystallographic refinement was performed with phenix.refine[29], including the addition of hydrogens in riding positions and TLS refinement. 5% of random reflections were flagged for the calculation of Rfree. The reported resolution limit of 2.67 Å was determined by paired refinement via the PDB_REDO server[30], and the dataset was reprocessed employing this high-resolution cutoff. The structural model was refined to R/Rfree of 21/24% in space group I222. The coordinates have been deposited in the PDB with the identifier 29ST. Data collection and refinement statistics are summarised in Table S14. Figures of crystal structures were prepared using the PyMOL Molecular Graphics System version 3.1.6 (Schrödinger, LLC).

Table S14: X-ray data collection and refinement statistics.

| **Structure** | AzeJ from *Saccharothrix* sp. NRRL B-16348 |
|---|---|
| PDB-ID: | 29ST |
| **Data collection** | |
| Beamline | Beamline P11, PETRA III, DESY, Hamburg |
| Wavelength (Å) | 0.99990 |
| Space group | I222 |
| Cell dimensions | |
| $a$, $b$, $c$ (Å) | 16.30, 150.57, 194.62 |
| $\alpha$, $\beta$, $\gamma$ (°) | 90, 90, 90 |
| Resolution (Å)[a] | 46.08 – 2.67 (2.72 – 2.67) |
| $R_{merge}$ (%)[a] | 24.7 (356.3) |
| $R_{pim}$ (%)[a] | 4.8 (69.9) |
| Unique reflections[a] | 48599 (2328) |
| $I/\sigma I$[a] | 12.7 (1.0) |
| Completeness (%)[a] | 99.9 (97.0) |
| Redundancy[a] | 27.4 (25.6) |
| $CC_{1/2}$ (%)[a] | 99.8 (59.7) |
| **Refinement** | |
| Resolution (Å) | 2.67 |
| No. reflections | 48554 |
| $R_{work/}$ $R_{free}$ (%) | 21.10/23.95 |
| No. atoms | 9042 |
| Protein | 9030 |
| Ligand | 0 |
| Water | 12 |
| B-factors (Å$^2$) | 74.27 |
| Protein | 74.29 |
| Ligand | - |
| Water | 57.29 |
| R.m.s deviations | |
| Bond lengths (Å) | 0.004 |
| Bond angles (º) | 0.641 |
| Ramachandran statistics (%) | |
| Favored | 97.24 |
| Allowed | 2.67 |
| Outliers | 0.10 |
| Clashscore (MolProbity) | 4.85 |
| MolProbity score | 1.39 |

[a]Values for the highest resolution shell are shown in parentheses.

##### *1.7.5 Crystal structure of SacAzeJ*

The crystal structure of *Sac*AzeJ was determined in the absence of ligand and deposited in the PDB under the identifier 29ST. Figure S22 shows a structural comparison with ligand-bound *Pa*AzeJ.

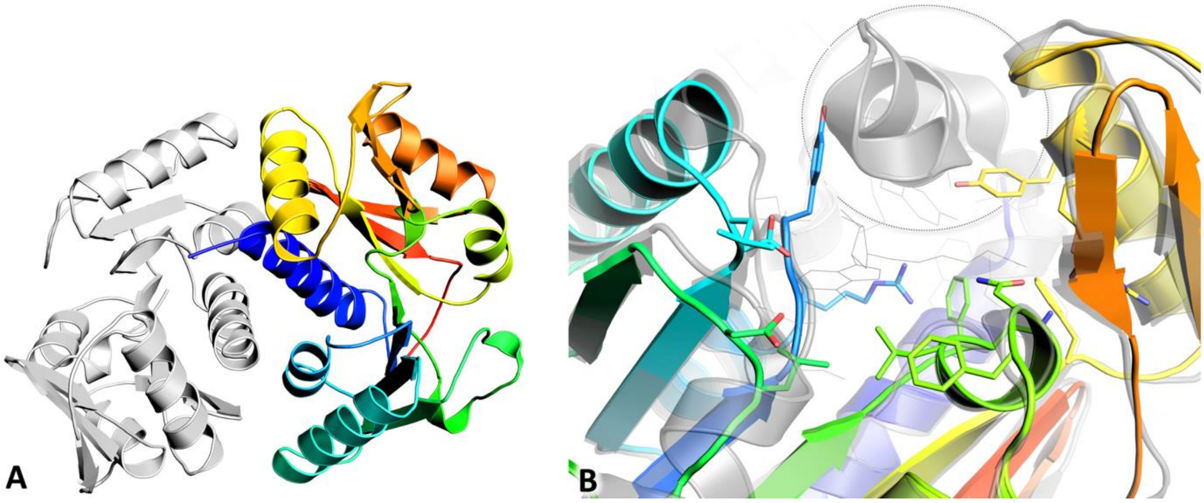

Figure S22: Crystal structure of AzeJ from *Saccharotrix* sp. NRRL B-16348 (SacAzeJ, PDB: 29ST) at 2.7 Å resolution. A: Dimer of SacAzeJ with one polypeptide chain colored from the N-terminus (blue) to the C-terminus (red). B: Magnification of the ligand binding site superimposed with AzeJ from *P. aeruginosa* (PaAzeJ, PDB: 8RYD, grey/transparent) with bound SAH shown as lines. Overall sequence identity is 60% with an alignment r.m.s.d. of 0.87 Å. Residues involved in SAM/SAH binding in PaAzeJ (grey lines) are conserved in SacAzeJ (sticks). The *N*-terminal helix (dashed circle) latching over the ligand binding site in the PaAzeJ ligand complex is not resolved in the apo SacAzeJ structure.

#### 1.8 Amino Acid Sequences of Enzymes

The underlined sequences represent the His-Tag or in the *Sac*AzeJ sequence the Strep-Tag.

***Uu*MAT:**

MQYKKIITSESVGAGHPDKICDQISDAILDECLSQDQNSRVACEVLACNRLIVIAGEITTHAYVDVVKTA
WEIIKPLGYDENDFTIISNVNKQSVDIAQSVDKTNKNLIGAGDQGIVFGYACDETPQYMPLTSVLAHELL
KEIERQRRSKEFIKIQADMKSQVSIDYSNSTPLIETMLVSIQHDEDYDVEYFNKKVSAIMEQIAKKYNLN
TNFKKIINSSGRFVIGGPIGDTGLTGRKIIVDTYGGVGHHGGGAFSGKDPTKVDRSASYFARWIAKNVV
AAKLAKQCEIQLAFAIGQPQPVAMYVNTFNTNLIDETKIFEAIKKSFNFDIKTFINDLNLWTTKYLPVATY
GHFGRDDLDLSWEKLNKVEDLIKNSKHHHHHH

***Tk*MAT:**

MGSSHHHHHHSSGLVPRGSHMAGKVRNIVVEELVRTPVEMQKVELVERKGIGHPDSIADGIAEAVSR
ALSREYVKRYGIILHHNTDQVEVVGGRAYPQFGGGEVIKPIYILLSGRAVEMVDREFFPVHEIALKAAK
DYLRKAVRHLDLEHHVIIDSRIGQGSVDLVGVFNKAKKNPIPLANDTSFGVGYAPLSETEKIVLETEKYL
NSDEFKKKYPAVGEDIKVMGLRKGDEIDLTIAAAIVDSEVDNPDDYMAVKEAIYEAAKGIVESHTERPT
NIYVNTADDPKEGIYYITVTGTSAEAGDDGSVGRGNRVNGLITPNRHMSMEAAAGKNPVSHVGKIYNI
LSMLIANDIAEQVEGVEEVYVRILSQIGKPIDEPLVASVQIIPKKGYSIDVLQKPAYEIADEWLANITKIQK
MILEDKVNVF

**CouO:**

MGSSHHHHHHSSGLVPRGSHMKIEPITGSEAEAFHRMGSRAFERYNEFVDLLVGAGIADGQTVVDLC
CGSGELEIILTSRFPSLNLVGVDLSEDMVRIARDYAAEQGKELEFRHGDAQSPAGMEDLLGKADLVVS
RHAFHRLTRLPAGFDTMLRLVKPGGILNVSFLHLSDFDEPGFRTWVRFLKERPWDAEMQVAWALAH
YYAPRLQDYRDALAQAADETPVSEQRIW VDDQGYGVATVKCFARRAAA

***Pp*CaOMT:**

MGSSHHHHHHSSGLVPRGSHMASSLERKSHPKINHAEPEDEITKEEEDESFCYAMQLVGSSVLSMSL
QSAIKLGIFDIIARKGPGAKLSSSEIATKIGTENPEAPVMVDRILRLLTSHSVLNCSAVAANGGSDFQRV
YSLGPVSKYFVNDEEGGSLGPLLTLIQDRVFLESWSQLKDAVVEGGIPFNRVHGMHAFEYPGLDPRF
NQVFNTAMFNHTTIVIKKLLHIYKGLEDKNLTQLVDVGGGLGVTLNLITSRYQHIKGINFDLPHVVNHAP
SYPGVEHVGGDMFASVPSGDAIFMKWILHDWSDEHCLKLLKNCYKAIPDNGKVIVVEALLPAMPETST
ATKTTSQLDVLMMTQNPGGKERSEQEFMALATGAGFSGIRYECFVCNFWVMEFFK

***Rg*ANMT:**

MGSSHHHHHHSSGLVPRGSHMGSLSESHTQYKHGVEVEEDEEESYSRAMQLSMAIVLPMATQSAIQ
LGVFEIIAKAPGGRLSASEIATILQAQNPKAPVMLDRMLRLLVSHRVLDCSVSGPAGERLYGLTSVSKY
FVPDQDGASLGNFMALPLDKVFMESWMGVKGAVMEGGIPFNRVHGMHIFEYASSNSKFSDTYHRAM
FNHSTIALKRILEHYKGFENVTKLVDVGGGLGVTLSMIASKYPHIQAINFDLPHVVQDAASYPGVEHVG
GNMFESVPEGDAILMKWILHCWDDEQCLRILKNCYKATPENGKVIVMNSVVPETPEVSSSARETSLLD
VLLMTRDGGGRERTQKEFTELAIGAGFKGINFACCVCNLHIMEFFK

***Ec*MTAN:**

MGSSHHHHHHSSGLVPRGSHMKIGIIGAMEEEVTLLRDKIENRQTISLGGCEIYTGQLNGTEVALLKSG
IGKVAAALGATLLLEHCKPDVIINTGSAGGLAPTLKVGDIVVSDEARYHDADVTAFGYEYGQLPGCPAG
FKADDKLIAAAEACIAELNLNAVRGLIVSGDAFINGSVGLAKIRHNFPQAIAVEMEATAIAHVCHNFNVPF
VVVRAISDVADQQSHLSFDEFLAVAAKQSSLMVESLVQKLAHG

**SgvM:**

MGSSHHHHHHSSGLVPRGSMATHDIAAQHLADGIAASGPAPDLAAAAAFLEMGDRLGVVAHLDPDR
TLETAEVAAALDLPEPALVRYLDAVESAGLVIREGEGRYRACPDFDTIRHQAGYISWTMNANRPFIENA
RDFFTDWDKAARTHVRDYREVAVSSQWMGSHAFYPTALATIIDAAPRKVVDLGAGTCRLLIEVLGAVP
GSTGVGLDFAADACRAAEQAVAQAGMTDRLTVVERTIQSVATDPGVLEGADVIHAGFVFHDMLPEEE
DVCDQVLANCRESLAPGGFLAITDAVPYLRNDRERRFSAAVSYYHGEFMRRRLQSEEEWVERLRGA
GFSDVRALTLAFPTGRLFLAHR

***Pa*AzeJ:**

MGSSHHHHHHSSGVRGSHMSQNMDLTIGTHRESAFYELEFGPRTIMTLANFPDDVLPLLQMESLMTF
EAMAYLRCDALVELGCYDGRALEIARLLNARYLGVDLDQRAIETLRTRIEREGMSDRADTVVDDILNHT
RRGASVGSRALYLLPFNLLGNFREPKRLLDSLAERSVAAVVSVFGDSAEATRVRQSYYRRCGVQGLE
LHTRDDGTVFTGSDGFYSRSYSRACLHALLAECGLTVVRSASNLFAHCVTVLPEGADQGFGSSAA

***Sac*AzeJ**

MASWSHPQFEKVDENLYFQGGGRAVEHQPESAFYARSSGSATIMTLANFPDDVLPLLQMESLATVE
AMPYLGCDTLVELGCYDGRALEVARHLGVRYLGVDLDARAVATLAARIEREGMADRARTLVDDLFNH
HRWDRDVVGERPLYMLPFNLLGNFRHPLRLLRSLAGASGAAVISVFGEGLEATKVRHAYYTRCGVRG
LEFELTGDGGVLFTGADGFYSRSFSESRFRALLRECGLTVVRVRGNSLGYAATVLLGDAPSEARRR

**GenD1**

MGSSHHHHHHSQDPMTVTNKIVTGVAFPPSLLAETPPISVATLTAYLRDKGMPAVGLDLNADFNEYLL
NRVEIEQVQGPENTHEFTKPFIKQFFLNHITGNYFTETNFEQWDLQQQCQVAPESLSIWDPPFPFSYC
EFLSILRDEPERVAKLVRDPDANIYHAFYQEKVAGKASELGLMGFSIMGYNQVIPALTLGYLMKKENPD
LYICWGGPWVTSFADMLIPRLEACPELGELIDALVVREGEEPLLKMAEALSRGERPVGIPGGCKPMSE
QSVLDTKPVGRKLLLAMSQPRENMPNTHWRIEDGTYERSGDISWVADMNQLPTPDYSDFDLSLYTTF
REGQGSLVLQGSRSCYYMKCSFCNAITNFAPWSYRERSTENIQKDIDTFLELYPGTVHFDFADAVFPA
KRLVELADFFIAKKRPELFWEVDVRFEGNIDKAVLTKMRDSQGTLRFGLETANERLLDLVRKGNRMEV
VHRLLQDSRELGYKPFLMTIVGLPTEEREEAEELYQFLSDYHDTVTYQIADFIVERNSPIQLRPDDYGIH
IDDDEQESFHHNLHFTRRAGYSDEEAQEVYRDILVRTMQRFKGAHEVDVEQERSRVAPDDSVYRLSL
RAGSFALENYWVKHNNLPFEGLVPIGYKVQQQWTDMTDKGTVFEIDPDIALGALAGAGSR

**HydG**

MGSSHHHHHHSQDPMVKEKADFINDEKIRQDLEKAKKATSKDALEIIEKAKNLKGITPEEAAVLLNVED
EDLLNEMFKVARYIKEEIYGNRIVIFAPLYVSNYCVNNCRYCGYRHSNEQQRKKLTMEEVRREVEILEE
MGHKRLAVEAGEDPVNCPIDYIVDVIKTIYDTKLKNGSIRRVNVNIAATTVENYKKLKKVGIGTYVLFQE
TYHRPTYEYMHPQGPKHDYDYHLTAMDRAMEAGIDDVGLGVLYGLYDYKYETVAMLYHANHLEEKF
GVGPHTISVPRLRPALNISIDKFPYIVSDKDFKKLVAVIRMAVPYTGMILSTREKPKFREEVISIGISQISA
GSCTGVGGYHEEISKKGGSKPQFEVEDKRSPNEILRTLCEQGYLPSYCTACYRMGRTGDRFMSFAK
SGQIHNFCLPNAILTFKEFLIDYGDEKTKKIGEKAIAVNLEKIPSRTVREETKRRLTRIENGERDLYF

**QCMT**

MDITIFSPGIYTYGAMLIGGVLRDAGHEVTLTRTPEVPEGSLLLASLFSTQHLLDPKIRSLVRRHRERGG
TVYVGGPVSAVPEIVLRELAPDAVVVGEGEETVVRLAEEGASETLPGIAYLDGDQAVVTAPAPPAPIDR
PLPLIPEDIGSQSIRGASAYIETHRGCIGGCTFCQVPRFFGREVRSRPLESIIEEVKAFRAAGAKRLSISG
GTGSLYGSHGCEMNPGAFIALLRGMAEVMGPKNVSAPDIRVDCITDEILDAIRQYTIGWVFFGIESGSN
RVLRQMGKGATVREVEEAVERCREHGLHVAGSFIVGYPGETERDYEATKDLVAALSLDDVFISSAEPI
PGTPLADLVIRTPRDKNPAFMPHTGEYRALHLTESEARCFDLMMHADMYRPQIRLVTDEVYAAYLAEA
KKQGEDIRAATELILRYAQRLENLYFQGHHHHHH

**TsrM**

MGSSHHHHHHSQDPMLRKGTVALINPNQIHPPIAPYALDVLTTALEASGFEAHVLDLTFHLDDWRQTL
RDYFRAERPLLVGVTCRNTDTVYALEQRPFVDGYKAVIDEVRRLTAAPVVAGGVGFSTMPFALVDYF
GIEYGVKGPGEKIICDLARALAEGRSADRIHIPGLLVNRGPGNVTRVAPPALDPRAAPAPSSSPSPSPA
PSSSSAPVPVPLSFAAVGHHESRAWQAETELPYTRRSGEPYKVDNLRYYREGGLGSILTKNGCVYKC
SFCVEPDAKGTQFARRGITAVVDEMEALTAQGIHDLHTTDSEFNLSIAHSKNLLREIVRRRDHDATSPL
RDLRLWVYCQPSPFDEEFAELLAAAGCAGVNIGADHTRPEMLDGWKVTAKGTRYYDFADTERLVQL
CHRNGMLTMVEALFGMPGETLETMRDCVDRMMELDATVTGFSLGLRLLPYMGLAKSLAEQCDGVRT
VRGLQSNNASGPIVLKQLHQCDGPIEYERQFMFDESGDFRLVCYFSPDLPEAPGTADSPDGIWRASV
DFLWDRIPKSEQYRVMLPTLSGSSENDNNYADNPFLTSLNRKGYTGAFWAHWRDREAIMSGATLPL
GELAEAVR

**ACCS**

HHHHHHSSGGENLYFQGHMRMLSRNATFNSHGQDSSYFLGWQEYEKNPYHEVHNTNGIIQMGLAE
NQLCFDLLESWLAKNPEAAAFKKNGESIFAELALFQDYHGLPAFKKAMVDFMAEIRGNKVTFDPNHLV
LTAGATSANETFIFCLADPGEAVLIPTPYYPGFDRDLKWRTGVEIVPIHCTSSNGFQITETALEEAYQEA
EKRNLRVKGVLVTNPSNPLGTTMTRNELYLLLSFVEDKGIHLISDEIYSGTAFSSPSFISVMEVLKDRNC
DENSEVWQRVHVVYSLSKDLGLPGFRVGAIYSNDDMVVAAATKMSSFGLVSSQTQHLLSAMLSDKKL
TKNYIAENHKRLKQRQKKLVSGLQKSGISCLNGNAGLFCWVDMRHLLRSNTFEAEMELWKKIVYEVH
LNISPGSSCHCTEPGWFRVCFANLPERTLDLAMQRLKAFVGEYYNVPEVNGGSQSSHLSHSRRQSL
TKWVSRLSFDDRGPIPGR

#### 2. References

[1] a) E. Jockmann, F. Subrizi, M. K. F. Mohr, E. M. Carter, P. M. Hebecker, D. Popadić, H. C. Hailes, J. N. Andexer, *ChemCatChem* **2023**, *15*, e202300930; b) J. Siegrist, S. Aschwanden, S. Mordhorst, L. Thöny-Meyer, M. Richter, J. N. Andexer, *ChemBioChem* **2015**, *16*, 2576; c) S. Mordhorst, J. Siegrist, M. Müller, M. Richter, J. N. Andexer, *Angew. Chem. Int. Ed.* **2017**, *56*, 4037; d) F. Michailidou, N. Klöcker, N. V. Cornelissen, R. K. Singh, A. Peters, A. Ovcharenko, D. Kümmel, A. Rentmeister, *Angew. Chem. Int. Ed.* **2021**, *60*, 480; e) C. Sommer-Kamann, J. Breiltgens, Z. Zou, S. Gerhardt, R. Saleem-Batcha, F. Kemper, O. Einsle, J. N. Andexer, M. Müller, *ChemBioChem* **2024**, *25*, e202400258; f) Z. Hong, A. Bolard, C. Giraud, S. Prévost, G. Genta-Jouve, C. Deregnaucourt, S. Häussler, K. Jeannot, Y. Li, *Angew. Chem. Int. Ed.* **2019**, *58*, 3178; g) D. Kleiner, F. Shmulevich, R. Zarivach, A. Shahar, M. Sharon, G. Ben-Nissan, S. Bershtein, *Journal of Molecular Biology* **2019**, *431*, 4796.

[2] J. Gagsteiger, S. Jahn, L. Heidinger, L. Gericke, J. N. Andexer, T. Friedrich, C. Loenarz, G. Layer, *Angew. Chem. Int. Ed.* **2022**, *61*, e202204198.

[3] P. Dinis, D. L. M. Suess, S. J. Fox, J. E. Harmer, R. C. Driesener, L. de La Paz, J. R. Swartz, J. W. Essex, R. D. Britt, P. L. Roach, *Proceedings of the National Academy of Sciences of the United States of America* **2015**, *112*, 1362.

[4] N. D. Lanz, A. J. Blaszczyk, E. L. McCarthy, B. Wang, R. X. Wang, B. S. Jones, S. J. Booker **2018**, *57*, 1475.

[5] P. Montenegro, I. M. Valente, L. M. Gonçalves, J. A. Rodrigues, A. A. Barros, *Anal. Methods* **2011**, *3*, 1207.

[6] Jana Gagsteiger, *Dissertation*, University of Freiburg, **2023**.

[7] a) C. Huang, F. Huang, E. Moison, J. Guo, X. Jian, X. Duan, Z. Deng, P. F. Leadlay, Y. Sun, *Chem. Biol.* **2015**, *22*, 251; b) S. Pierre, A. Guillot, A. Benjdia, C. Sandström, P. Langella, O. Berteau **2012**, *8*, 957.

[8] D. Moreno-González, F. J. Lara, N. Jurgovská, L. Gámiz-Gracia, A. M. García-Campaña, *Analytica Chimica Acta* **2015**, *891*, 321.

[9] F. Yan, R. Müller, *ACS chemical biology* **2019**, *14*, 99.

[10] https://www.rcsb.org/pages/policies.
[11] J. Boitreaud, J. Dent, M. McPartlon, J. Meier, V. Reis, A. Rogozhnikov, K. Wu, *Chai-1: Decoding the molecular interactions of life*, **2024**.
[12] X. Tan et al., *Worldwide Protein Data Bank*, **2008**.
[13] https://www.pymol.org/support.html.
[14] T. Pavkov-Keller, K. Steiner, M. Faber, M. Tengg, H. Schwab, M. Gruber-Khadjawi, K. Gruber, *PloS one* **2017**, *12*, e0171056.
[15] S. Ju, K. P. Kuzelka, R. Guo, B. Krohn-Hansen, J. Wu, S. K. Nair, Y. Yang, *Nat. Commun.* **2023**, *14*, 5704.
[16] T. J. Klaubert, J. Gellner, C. Bernard, J. Effert, C. Lombard, V. R. I. Kaila, H. B. Bode, Y. Li, M. Groll, *Nat. Commun.* **2025**, *16*, 1348.
[17] G. Capitani, A. C. Eliot, H. Gut, R. M. Khomutov, J. F. Kirsch, M. G. Grütter, *Biochim. Biophys. Acta* **2003**, *1647*, 55.
[18] J. L. Galman, F. Parmeggiani, L. Seibt, W. R. Birmingham, N. J. Turner, *Angew. Chem. Int. Ed.* **2022**, *61*, e202112855.
[19] K. D. Nakamura, F. Schlenk, *Arch. Biochem. Biophys.* **1976**, *177*, 170.
[20] F. W. Studier, *Protein Expression and Purification* **2005**, *41*, 207.
[21] A. Burkhardt, T. Pakendorf, B. Reime, J. Meyer, P. Fischer, N. Stübe, S. Panneerselvam, O. Lorbeer, K. Stachnik, M. Warmer et al., *Eur. Phys. J. Plus* **2016**, *131*, 56.
[22] C. Vonrhein, C. Flensburg, P. Keller, A. Sharff, O. Smart, W. Paciorek, T. Womack, G. Bricogne, *Acta crystallographica. Section D, Biological crystallography* **2011**, *67*, 293.
[23] W. Kabsch, *Acta crystallographica. Section D, Biological crystallography* **2010**, *66*, 125.
[24] P. Evans, *Acta Cryst D* **2006**, *62*, 72.
[25] P. R. Evans, G. N. Murshudov, *Acta crystallographica. Section D, Biological crystallography* **2013**, *69*, 1204.
[26] A. J. McCoy, R. W. Grosse-Kunstleve, P. D. Adams, M. D. Winn, L. C. Storoni, R. J. Read, *J Appl Cryst* **2007**, *40*, 658.
[27] Adams PD, *Acta Crystallogr D Biol Crystallogr* **2009**, *65*, 1074.
[28] P. Emsley, B. Lohkamp, W. G. Scott, K. Cowtan, *Acta crystallographica. Section D, Biological crystallography* **2010**, *66*, 486.
[29] P. V. Afonine, R. W. Grosse-Kunstleve, N. Echols, J. J. Headd, N. W. Moriarty, M. Mustyakimov, T. C. Terwilliger, A. Urzhumtsev, P. H. Zwart, P. D. Adams, *Acta crystallographica. Section D, Biological crystallography* **2012**, *68*, 352.
[30] R. P. Joosten, F. Long, G. N. Murshudov, A. Perrakis, *IUCrJ* **2014**, *1*, 213.
